# Sex differences in cognitive performance, style and domain relationships in mosquitofish (*Gambusia affinis*)

**DOI:** 10.1101/842278

**Authors:** Kelly J. Wallace, Richie T. Rausch, Mary E. Ramsey, Molly E. Cummings

**Affiliations:** Department of Integrative Biology, University of Texas, Austin, TX, 78712, USA

**Author notes:** **Author for correspondence**: Kelly J. Wallace, University of Texas at Austin, Department of Integrative Biology, 1 University Station C0990, Austin, Texas 78712.

**Keywords:** sex differences, cognitive style, numerical discrimination, cognitive flexibility, *poeciliidae*, *Gambusia affinis*

## Abstract

Given that the sexes often differ in their ecological and sexual selection pressures, sex differences in cognitive properties are likely. While research on sexually dimorphic cognition often focuses on performance, it commonly overlooks how sexes diverge across multiple cognitive tasks (cognitive domains) and in behaviors associated with cognitive performance (cognitive style). We tested male and female western mosquitofish (*Gambusia affinis*) in three cognitive tasks: associative learning (numerical discrimination), cognitive flexibility (detour task), and spatio-temporal learning (shuttlebox). We characterized statistical relationships between cognitive performances and cognitive style during the associative learning task with measures of anxiety, boldness, exploration, reaction time, and activity. We found sex differences in performance, cognitive style, and the relationships between cognitive domains. Females outperformed males in spatio-temporal learning task, while the sexes performed equally in associate learning and cognitive flexibility assays. Females (but not males) exhibited a ‘fast-exploratory’ cognitive style during associative learning trials. Meanwhile, only males showed a significant positive relationship between domains (associative learning and cognitive flexibility). We propose that these sexually dimorphic cognitive traits result from strong sexual conflict in this taxon; and emphasize the need to explore suites of sex-specific cognitive traits and broader comparative work examining sexual selection and cognition.

**Highlights:**

- Males and females perform at similar levels in associative learning and cognitive flexibility assays, but females tend to outperform males on a spatio-temporal learning task.
- Female performance in associative learning trials (numerical discrimination task) can be predicted by cognitive style behaviors (exploration, reaction time, and activity); whereas male performance cannot.
- Males, but not females, show a predictive relationship between associative learning and cognitive flexibility performance.
- Our results demonstrate that sex differences in cognition extend beyond performance into cognitive style and domain relationships, suggesting that investigations into animal personality and cognition require more comprehensive characterization.

## 1. Introduction

Individuals vary in many cognitive attributes such as learning ability, style, and consistency in performance across varying tasks. These three attributes commonly referred to as cognitive performance, cognitive style, and cognitive domain respectively, are often examined individually, but recent efforts have begun to identify how they may be inter-related [1–3]. For instance, some of the early work in this arena has suggested that how quickly an animal makes a decision and how much it explores its environment (e.g. a fast-exploring cognitive style) should predict an individual’s performance (accuracy) in a learning task [2]. Empirical studies have shown variable support for this hypothesis with some taxon revealing a positive relationship [4] between fast-exploratory styles and learning performance, others revealing a negative relationship [5–6], and yet other studies find that a fast-exploratory cognitive style is unrelated to performance [7]. Furthermore, relationships between cognitive style and performance are often domain-specific [8]. Understanding how and why these relationships vary across taxa and between domains is a current challenge in cognitive studies, however, a factor that is emerging as one of the predominant predictors of variation across cognition is sex [8].

A recent meta-analysis by Dougherty & Guillette (2018) identified sex as the single best categorical variable that explains the variation across animal studies on the associations between personality and cognition [8]. This is unsurprising given fitness benefits for particular cognitive traits are often sex-dependent [9–13], leading to differential selective pressures favoring certain cognitive traits in one sex and potentiating sexual dimorphism. While cognitive sex differences are accumulating across the literature, there exists a great deal of variation between and within taxa [14] and very little work examining how these sex differences bear out across cognitive domains. Sex differences in cognitive performance are frequently domain specific [15–21], and even when learning performance is equivalent between the sexes the behavioral predictors that underlie individual variation can differ [22–23]. The present study is designed to specifically examine sex differences in cognitive performance, cognitive style, and relationships between cognitive domains in a taxon with well-defined sexually dimorphic behaviors [24–27].

The *poeciliidae* family of livebearing freshwater fish— which includes the guppies, mollies, mosquitofish, and swordtails— exhibits a wealth of natural variation in sexual selection pressures [24] and cognition [25], making this family a uniquely suitable system to address sex differences in cognition [26]. Poeciliids are famous for sexually divergent reproductive roles as males provide only sperm via an intromittent organ (gonopodium) and females undergo a monthlong internal gestation period. Mosquitofish (*Gambusia sp*.) are characterized by a high degree of sexual conflict, with males exhibiting some of the highest rates of sexual harassment across the poeciliidae family and females adopting a strong avoidance responses including shoaling to reduce male harassment [27]. Artificial selection experiments on male gonopodial length in this genus results in larger female (but not male) brain size [28], suggesting that sexual conflict may act differentially on cognition. In our study we assessed the cognitive performance of the western mosquitofish (*Gambusia affinis*) in three cognitive domains: associative learning (numerical discrimination assay), cognitive flexibility (detour task), and spatio-temporal learning (shuttlebox assay). To evaluate cognitive style, we measured the relationship between associative learning performance and a variety of behaviors displayed during the numerosity discrimination test trials. We then examined whether performance in one cognitive domain predicted performance in another and whether these relationships varied by sex.

By examining sex differences in cognitive performance across associative learning, cognitive flexibility and spatio-temporal learning, we can begin to determine how divergent sexual selection pressures influence these cognitive domains. Based on previous studies with this species, we expect similar numerical discrimination performances between the sexes [22]. Based on work with guppies, we predict that females will outperform males in a detour task [15,17]. Our spatio-temporal assay (shuttlebox) has only previously been performed on one other fish taxon (zebrafish [29]) with no sex-dependent effect. However, given the strong selection pressures on females to find refuge from male harassment, we predicted that female *G. affinis* are likely to outperform male *G. affinis* in a spatio-temporal learning assay that varies time and place of a shoal group. Moreover, we predicted that the sexes would diverge in cognitive style where a ‘fast-exploratory’ learning type (e.g. faster decision making, shorter latencies to sample) would be more associated with males as has been found in other poeciliids [7]. Lastly, we predicted that the sexes would diverge in their relationship between cognitive domains. While the literature generally suggests a negative relationship between associative learning and cognitive flexibility [30], studies in poeciliids have thus far have not found a relationship [15,31]. Domain relationships between spatio-temporal learning and associative learning have not yet been explored in poeciliids, however, we hypothesized that *Gambusia affinis* females will exhibit a positive relationship between these two domains as both are critical to shoaling decisions.

## 2. Materials and Methods

### (a) Housing

Wild-caught western mosquitofish *Gambusia affinis* (27 female, 27 male) from outdoor ponds at Brackenridge Field Laboratories in Austin, TX were group housed in 35 gallon aquaria at 24.4-26.6◻C on a 13-11 light cycle. Prior to testing, individuals were socially isolated for 2 days in 2.5 gallon aquaria. Individuals participated in 13 days of cognitive assay testing with assay order balanced across individuals and sexes, and 24 hour interval between assays.

### (b) Numerical Discrimination Experimental Design

To test associative learning individuals were placed in a modified 10-gallon automated numerical discrimination experimental tank for an 11 day assay including 2 days of habituation, 6 days of training, and 3 days of testing (Supplementary Fig 1). Stimuli were geometric shapes (adapted from Etheredge et al 2018 and controlled for non-numerical cues) [22] presented on LCD screens attached to the end sides of each tank. Version control of the automation scripts are available at https://github.com/jenkins-cummingslab/ethoStim; and for a more detailed description of our automated numerical discrimination operation see *Supplementary Methods*.

Trials occurred five times daily during habituation and training. During habituation, a food reward was administered simultaneously at both blank screens for four minutes. During training individuals were presented two 1:2 ratios (5 versus 10 shapes or 6 versus 12 shapes) haphazardly alternating between left and right reward sides across training trails. Stimuli appeared on screens for an initial 10 seconds prior to a food reward descending into the tank for an additional 10 seconds. Half of the individuals received food reward on the side with the greater quantity on the screen, and half of the individuals received food reward on the lesser quantity side. Individuals were then tested (no reward) for three days three times daily on novel testing ratios of varying difficulties: 1:2 (7 vs 14 shapes), 2:3 (8 vs 12 shapes), and 3:4 (9 vs 12 shapes), with reinforcement trials with original training ratios following each test trial to prevent extinction).

We assessed learning performance during the initial 20 seconds in which stimuli were presented on the screens which corresponded to the reward administration time period during training. Similar to other numerical discrimination studies [22,32], we employed a ‘learning criterion’ of individuals that had at least one testing ratio with a median performance value (proportion time spent in the closest quarter of the tank to the correct screen) above 60%.

### (c) Cognitive Flexibility (Detour maze) Experimental Design

The experimental tank (filled to 13cm) was subdivided into a starting alley (26 × 14cm), center section (14 × 31cm), and reward section (14 × 31cm). Because female *Gambusia affinis* shoal with other females to reduce male harassment [27,33] we used a male social activator and a female social reward. The male was placed behind the starting alley in a visible container. The center section contained a 25cm wide glass barrier which prevented individuals travelling in a direct line from reaching the social reward. The reward section contained the female in a visible container. Individuals could solve the detour task by turning away from the direct line and travelling through the unobstructed zones (3cm) at each side of the glass barrier to reach the social reward (Supplementary Fig 2). The focal individual habituated for five minutes in an opaque tube then swam freely for 10 minutes. Motivation was recorded as the latency to reach the barrier, and solution speed was recorded as the time difference between arrival at the barrier and reaching the social reward.

### (d) Temporal learning (Shuttlebox) Experimental Design

Our experimental tank was 52 × 26cm filled a depth of 10cm, with an Adafruit 7” LCD Display at either end. The focal individual swam freely during a 5-minute habituation in which both screens displayed a video of an empty tank. After habituation, one screen presented a 20 second video of 5 conspecific females, followed by a 90 second inter-stimulus interval (ISI) of the empty tank on both screens, then by the conspecific video on the opposite side (Supplementary Table 1). This alternating stimuli-ISI pattern lasted one hour as in [29].

Learners were evaluated as those who spent >50% of interaction time (within 10cm of screens) in the correct region (screen with an imminent shoal group appearing) during the final 30 seconds of the ISI for three consecutive trials following the 4^th^ ISI. Individuals who became non-active before the fourth ISI were not included in the analysis.

### (e) Video Scoring

Human observers scored time spent in regions of the numerical discrimination tank (using CowLog 3.0.2 and a python-generated grid overlay https://github.com/kjw2539/make_a_grid_eagle.py). Additionally, we recorded latency to change regions following the image presentation (reaction time), number of unique zones visited (exploration), and total transits between zones (activity). One author (KW) and twelve undergraduate students independently scored 315 videos (Single Score Intraclass Correlation between eleven scorers p = 5.41 × 10^−90^). Recordings for the Detour Maze and Shuttlebox assays were taken using Debut Video Capture Software and LifeCam cameras. Detour videos were independently scored by hand by two undergraduate student scorers and compared to the co-author (KW) scorer (p = 4.83 10^−45^ and p = 2.03 × 10^−35^). Shuttlebox videos were hand scored by three undergraduate student scorers using a python program developed by Luke Reding (see https://github.com/lukereding/shuttlebox/blob/master/track/shuttlebox_hand_track.py) that evaluated the position of the fish at 5 second intervals.

### (f) Statistics

We used multiple linear regressions to determine correlations between continuous variables. We conducted an unpaired t-test or unpaired Wilcoxon signed-rank test (determined by a Shapiro-Wilk normality test) for continuous data split into 2 categories, or a One-way ANOVA or Kruskal-Wallis test depending on normality for continuous data split into >2 categories. For categorical data we conducted Chi-squared tests on data with >5 observations per category and Fisher’s Exact Test for data which had <5 observations in a category. Data analysis and visualization were conducted using RStudio (3.2.2). Data analysis coding scripts, original data, and protocols can be found at https://github.com/kjw2539/Comparative-Cognition-R-scripts).

## 3. Results

54 individual *G. affinis* were run through a series of three cognitive assays (numerosity discrimination, detour maze, and shuttlebox), however, due to technical errors not all individuals completed all assays. Completion tallies across assays varied with 36 individuals completing numerosity discrimination (18 males, 18 females); 52 individuals completing detour maze (25 males, 27 females), and 26 individuals completing shuttlebox (9 males, 17 females).

### (a) Associative Learning Performance (Numerical Discrimination)

In the numerical discrimination task, 23 individuals successfully met the learning criterion (12F/11M) and 13 individuals did not (6F/7M). Sex did not influence the distribution of learners to non-learners (χ^2^ = 0; p = 1.00, Fig 1a). Our metric of performance correlated significantly with several others measured (Supplementary Fig 3; e.g. first side chosen (r^2^ = 0.175, p = 0.013), and latency to enter the correct side (r^2^ = 0.349, p = 0.0004)). There was no difference in performance between individuals trained to the higher quantity or lower quantity (p = 0.465) or across ratios (F = 0.302, p = 0.7399). The sexes did not differ significantly in their learning performance (t = 1.483, p = 0.148, Supplementary Fig 5). Body size (standard length), a proxy for age in female *G. affinis* [34], did not predict learning performance (females: p = 0.719, males: p = 0.770, combined: p = 0.589, see Table 1) and average standard length did not differ between individuals classified as learners and non-learners (females: p = 0.494, males: p = 0.285, combined: p = 0.839, Table 1).

**Fig 1.**
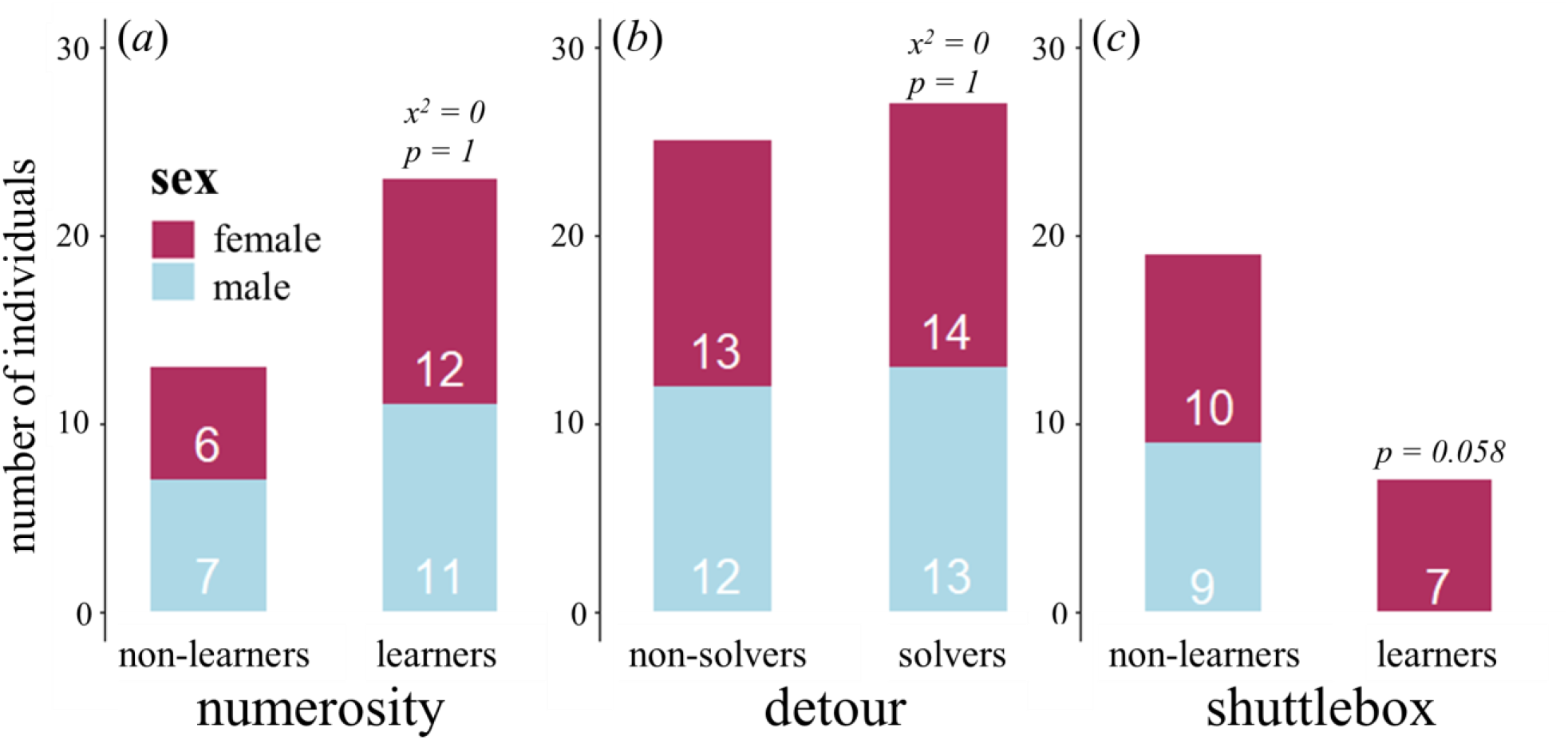
Learning performance differs between the sexes in the shuttlebox assay, but not the numerical discrimination or detour tasks. In the numerical discrimination assay, the sexes show equal distributions of learners and non-learners (learners reaching a minimum median performance of 60% for any of the 3 test ratios (7:14, 8 vs 12, 9 vs 12)) **(a)**. Detour maze solvers versus non-solvers also show equal distributions across the sexes **(b)**. Females reach the learning criterion in the shuttlebox assay (learners exhibiting three consecutive trials in which a majority of interaction time during the last 30 seconds of the ISI was spent within 10cm of the correct screen) more than males (Fisher’s Exact Test, p = 0.058) **(c)**.

**Table 1.** Female size predicts cognitive style behaviors in the numerical discrimination task. The relationship between standard length (a proxy for age) and the behavior listed is shown for three datasets: only females, only males, and all individuals combined. Continuous behavioral data results were determined using a multiple linear regression, and the reported effect size is Pearson’s correlation coefficient. Categorical behavioral data results (denoted with *) were determined using a t-test, and the reported effect size is Cohen’s D. Significant p-values are highlighted in red.

| Behavior | females<br>p | females<br>effect<br>size | males<br>p | males<br>effect size | both<br>sexes<br>p | both<br>sexes<br>effect<br>size |
| --- | --- | --- | --- | --- | --- | --- |
| Numerical Discrimination Performance | 0.719 | -0.094 | 0.770 | -0.074 | 0.589 | -0.095 |
| Numerical Discrimination Learning<br>Category* | 0.687 | -0.204 | 0.595 | 0.267 | 0.839 | -0.068 |
| Detour Solve Category* | 0.641 | 0.225 | 0.139 | 0.770 | 0.284 | 0.364 |
| Detour Latency to Solve | 0.344 | -0.358 | 0.219 | -0.454 | 0.537 | -0.156 |
| Detour Motivation | 0.438 | -0.217 | 0.455 | 0.218 | 0.795 | -0.050 |
| Shuttlebox Learning Category* | 0.434 | 6.676 | *na* | *na* | **0.034** | 5.504 |
| Numerical Discrimination Exploration | **0.021** | 0.539 | 0.182 | 0.329 | **0.009** | 0.430 |
| Numerical Discrimination Boldness | **0.048** | -0.471 | 0.706 | -0.096 | 0.423 | -0.138 |
| Numerical Discrimination Anxiety | 0.234 | 0.295 | 0.733 | -0.086 | 0.434 | -0.135 |
| Numerical Discrimination Activity | 0.052 | 0.464 | 0.122 | 0.378 | **0.0008** | 0.535 |
| Numerical Discrimination Reaction Time | 0.144 | -0.358 | 0.148 | -0.355 | **0.024** | -0.376 |

### (b) Behavioral Differences in Associative Learning (Numerical discrimination)

In the numerical discrimination assay, males and females exhibited the same levels of exploration (W = 192, p = 0.350), reaction time (W = 121, p = 0.203), sociability (W = 202, p = 0.214), boldness (t = 1.5621, p = 0.128) and activity (W = 217.5, p = 0.082). Males exhibited a higher proportion of time in regions associated with anxiety (thigmotaxis) than females (W = 97, p = 0.040, Supplementary Fig 4). Female standard length was significantly positively correlated with exploration (r = 0.539, p=0.021) and higher activity (r = 0.464, p = 0.052), but negatively correlated with boldness (r= −0.471, p = 0.048, see Table 1). Male standard length showed no significant relationship to any of these behaviors (Table 1).

### (c) Cognitive Style in Associative Learning (Numerical Discrimination)

Males and females differed in their relationships between behavior and performance. Female learning performance was significantly correlated with reaction time (r = −0.541, p = 0.025, Fig 2a) and exploration (r = 0.510, p = 0.036, Fig 2b), and marginally significant with activity r = 0.477, (p = 0.053, Fig 2c). Male behaviors did not relate to performance (reaction time: p = 0.689, exploration: p = 0.332, activity: p = 0.489, Fig 2d-f).

**Fig 2.**
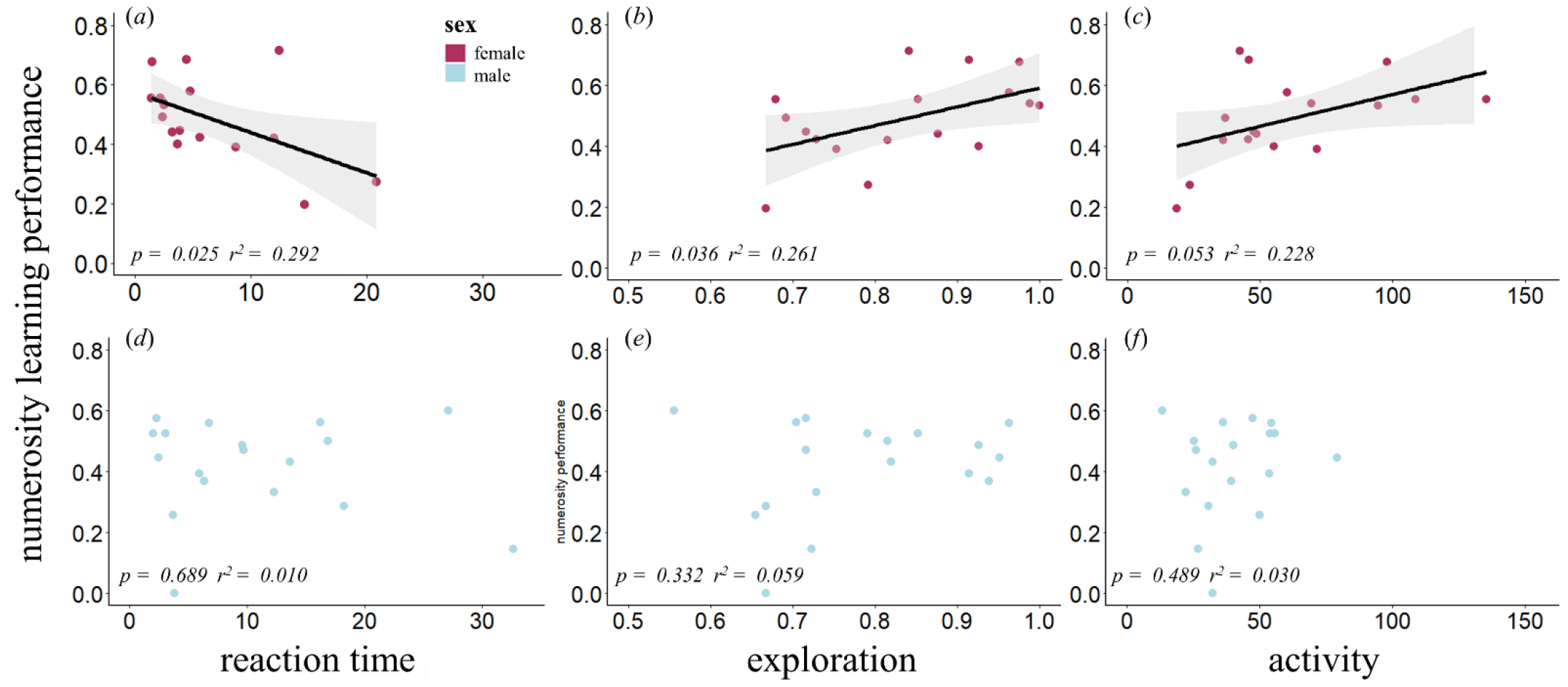
The sexes differ in cognitive style in the numerical discrimination task. Female learning performance (proportion time spent in contingency zone during initial 20 sec of test trial) is significantly predicted by female reaction time (p = 0.025)**(a)**, exploration (p = 0.036) **(b)**, and activity (marginal significance, p = 0.053) **(c)** displayed during test trials; whereas no significant correlations were found between these measures in males **(d-f)**.

### (d) Cognitive Flexibility Performance (Detour Maze)

In the detour maze, 27 individuals (14F/13M) successfully navigated around the transparent barrier, 25 did not (13F/12M) (Fig 1b). Of the 25 individuals who did not navigate around the barrier, 9 (4F/5M) did not participate (approach the barrier). Size did not differ between solvers and non-solvers (Table 1). On average it took solvers 80 seconds to solve the task upon reaching the barrier. Latency to solve the maze was not predicted by sex (W = 101, p = 0.645, Supplementary Fig 6), size (females: r = −0.358, p = 0.344, males: r = −0.454, p = 0.219, combined: r = −0.156, p =0.537), or motivation (latency to approach the barrier, r = −0.107, p = 0.596, Supplementary Fig 6). Motivation did not differ by sex (W = 221.5, p = 0.846) or size (females: r = −0.217, p = 0.438, males: r = 0.218, p = 0.455, combined: r = −0.050, p = 0.795) and did not influence whether an individual navigated around the barrier (W = 223.5, p = 0.860, Supplementary Fig 6).

### (e) Spatio-Temporal Learning Performance (Shuttlebox)

Of the 54 individuals tested in the shuttlebox assay, three were removed due to technical errors, and fifteen were removed from analysis for nonparticipation. We identified 7 learners (all female) and 19 non-learners(9M/10F). A near-significant sex difference in performance was found, where females reached the learning criterion more often than males (Fisher’s exact test, p = 0.058, Fig 1c).

### (f) Relationships in Performance across Domains

Males who solved the detour maze exhibited significantly higher numerical discrimination performance than non-solvers (t = −2.361, p = 0.035; Fig 3a). This relationship was not found in females (t= −0.673, p = 0.511).

**Fig 3.**
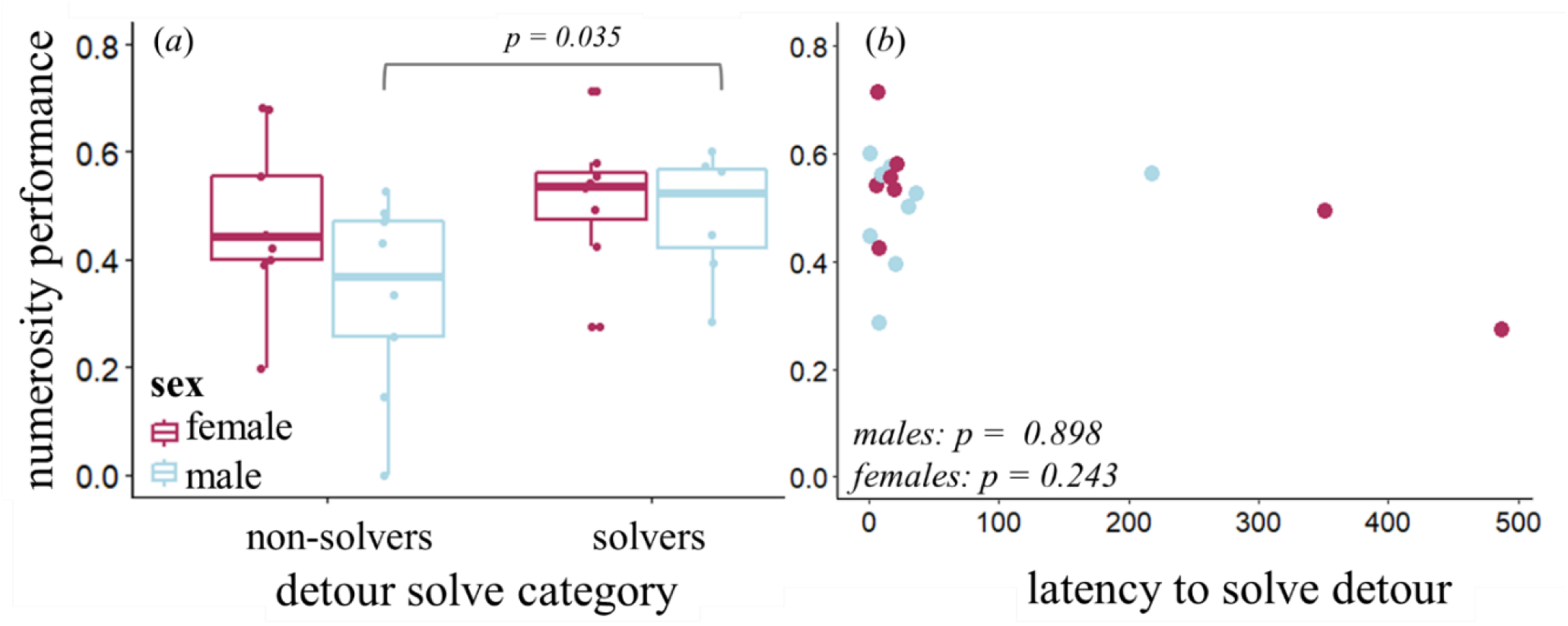
Male performance is predicted across cognitive domains. Males who solved the cognitive flexibility task (detour maze) exhibited significantly higher associative learning (numerical discrimination) performance than males who did not solve the cognitive flexibility task (p = 0.035) **(a)**. Latency to detour around the barrier once it was approached did not differ by sex and did not predict associative learning (numerical discrimination) performance **(b)**.

## 4. Discussion

We identified sex differences emerging across multiple attributes of cognition: in performance within a given task, in the cognitive styles that predict performance, and in the relationships in performance between cognitive tasks.

### (a) Sex Differences in Cognitive Performance

Similar to findings in previous poecilid studies utilizing numerical, color, and shape discrimination, we observed no sex differences in associative learning in our numerical discrimination assay [22,35]. We found similar ratios of learners to non-learners as other numerical discrimination experiments with *G. affinis* (two-thirds learners [22]). We found no sex difference in cognitive flexibility as measured via our detour task, which is contrary to findings in a related poecilid (*Poecilia reticulata*) where females outperform males in this task [15]. Confirming our prediction, we found a sex difference in performance in the spatio-temporal learning assay, with females being more likely to reach the learning criterion than males. The only previous examination of this cognitive task in teleosts showed no sex differences in zebrafish [29].

The greater spatio-temporal learning performance in female *G. affinis* may be driven by sexually dimorphic motivations to shoal. Females from poeciliid species with high levels of sexual coercion like *Gambusia* employ shoaling as a strategy to reduce male harassment[27], and *Gambusia* females shoal more than males [22,36]. Female *G. affinis* have been shown to choose shoal group size in a rational manner [37]; and the benefit for selecting larger shoals can lead to both reduced male harassment and increased foraging efficiency in a closely related poecilid (*Gambusia holbrooki*) [38]. Increased shoaling tendencies in females may drive the shoaling-related cognitive advantage observed in *G. affinis* females as seen in our spatio-temporal learning task in which females were more successful than males in predicting the time and place of the shoal. This sexual dimorphism in shoaling— and the sexual conflict that drives it— may manifest in sex differences in spatio-temporal cognitive tasks that emulate shoaling decisions.

### (b) Sex Differences in Cognitive Style

Performance in numerosity discrimination tasks was predicted by a suite of related behaviors exhibited during the test trials by females but not males. Female mosquitofish that exhibited a ‘fast-exploratory’ cognitive style in the numerical discrimination assay exhibited higher associative learning performance. Specifically, females that explored a greater area of the tank, reacted faster, and moved more demonstrated higher numerical discrimination performances. This ‘fast-exploratory’ type observed in these mosquitofish females appears to share attributes of previously described ‘fast behavioral’ type individuals (more exploratory, active, bold, and aggressive [2]) in other taxa. A positive association between ‘fast behavioral’ type and associative learning is documented across multiple taxa particularly in response to predation [8], including black-capped chickadees [439], sticklebacks [40], and Panamanian bishop fish *(Brachyraphis episcopi)* [41]. The ecological pressures that might lead to individuals adopting fast behavioral types may stem from being exposed to threatening environments (high predation), which places a selective pressure on the speed at which they sample their environment.

In some populations of guppies, intense male harassment has driven females into different habitats with greater predation levels than males [42]. Have the intense social pressures found in *Gambusia* driven sex-specific habitat differentiation and thus shaped female fast-exploratory cognitive styles? In previous fish studies examining sex differences in cognitive styles during associative learning assays with food rewards (T maze [43], visual discrimination of shapes and colors [44]) researchers have found a fast-exploratory cognitive style associated with *males*, not females. Only during shoal discrimination tasks in poeciliids (guppies, *Gambusia*) have females been shown to make faster decisions than males [7]. If strong sexual selection pressures have shaped female *Gambusia* shoaling decision-making processes, perhaps this has influenced the cognitive style in general numerical discrimination tasks.

The cognitive style that is identified in females during the numerosity discrimination task in this experiment is comprised of behaviors that tend to covary by female size (exploration, boldness and activity, Table 1). Poeciliid females experience indeterminant growth, and therefore standard length is often a proxy for female age. While larger/older females were no better at numerosity discrimination than smaller females, they did trend towards being more exploratory, reacting faster and moving more in the test trials (Table 1). This reflects findings showing that larger females tend to disperse farther [45]. Are female *G. affinis* developing a cognitive style as they age? Age-dependent decision-making processes have been documented among female poeciliids. For instance, relative to smaller females, larger and presumably older female El Abra swordtails show stronger preference for courting phenotypes over coercive phenotypes [46] and exhibit less transitivity in mate choice decisions involving male size [47]. Whether the increased exposure to male harassment over a female’s lifetime contributes to the development of fast-explorative female cognitive style can only be determined with manipulative social experiments.

It is imperative to caution that the causal relationship between cognitive style and cognitive performance is unknown [2,8]. In addition to implying that our ‘high learning individuals’ could be the result of individual differences in activity and exploratory tendencies, we must acknowledge that the behavior observed in the numerical discrimination testing trials could be a result of cognitive performance—i.e. those individuals who quickly learn the task may more quickly habituate and thus become more active and exploratory. Further studies assessing the developmental sources of variation in cognitive style and cognitive performance would help us understand the causal relationship between behavior and cognitive ability.

### (c) Sex Differences Across Cognitive Domains

In addition to performance and cognitive style, the sexes differed in their cross-domain relationships of performance: males (but not females) who successfully solved the cognitive flexibility task (detour maze) showed significantly higher associative learning performance. This relationship has not previously been described in poeciliid fish. Given that acquisition and reversal learning utilize different neural mechanisms [48–50], our finding of a positive relationship in male performance across these domains suggests non-modularity in male cognition, or a potential “g-factor” of general intelligence [51]. A general intelligence factor typically describes roughly 40% of individual variation in human tests, and g -factors are often found in studies across multiple animal taxa including birds, mice, chimpanzees, and dogs [14]. But what might explain our sex differences in the cross-domain relationships? *Gambusia* are extremely invasive (found in over 40 countries [52]) and thus frequently experience highly variable environments. They likely must utilize both associative learning *and* cognitive flexibility as a strategy to succeed in these dynamic environments. This concept, known as the “adaptive flexibility hypothesis,” emphasizes that cognitive flexibility is an adaptive response to a changing physical [53] or social [54] environment. An investment in a positive domain relationship between associative learning would predominantly benefit *Gambusia* males given that males are more likely to disperse than females [45] and disperse farther [55]. However, whether this sex-specific domain relationship is driven solely by potential ecological differences between the sexes or some contribution of different sexual selection pressures is yet to be determined.

## 4. Conclusion

Our study identified new sex differences in spatio-temporal learning, sex-specific cognitive styles in associative learning, and a sex-specific positive relationship between performance across cognitive domains. A wealth of literature has identified sex differences in cognitive performance in mammals [56], birds [18], reptiles [19], and fish [25–26]. But here we find that sex differences extend beyond performance into other cognitive attributes such as cognitive style and cross-domain relationship suggesting that more comprehensive characterization of cognition is important. Our experimental design in which the same individuals were assessed for cognitive performance and style across domains allowed us to find previously undescribed sex differences in cognition in *Gambusia affinis*. Fish exhibit a wealth of sex-specific ecological [20,57] and sexual selection pressures [24,26–27], therefore we can expect fish to continue to be an insightful taxonomic group in uncovering predictive patterns of sex differences in cognition [26]. Further studies, particularly those utilizing more extensive suites of cognitive testing, investigating neural mechanisms, and identifying developmental basis of these relationships will be critical to elucidate mechanisms governing the patterns observed in this study. In addition, comparisons of related species that differ in degree of sexual conflict and ecological pressures will be an important next step in distinguishing the factors that drive individual variation in cognitive performance.

## Supporting information

Electronic Supplementary Materials

## Data Accessibility

Raw data (Microsoft Excel) as well as analysis documentation are available at the following archived Github repository: https://github.com/kjw2539/Comparative-Cognition-R-scripts

## Acknowledgements

The authors would like to thank Luke Reding for contributions to the shuttlebox data analysis, the many undergraduates who assisted in data collection on this project including Matt Armstrong, Lauren Borland, Rahi Dakwala, Caleb Fleischer, Daniel Hauser, Amogh Kashyap, Presley Mackey, Claire Mayorga, Jessika McFarland, Lily Parsi, Huynh Pham, Sylvestre Pineau, Adam Redmer, Vishaal Sakthivelnathan, Eduardo Saucedo, Madison Schumm, Ben Whelan, Melody Ziari, and the very helpful comments and assistance from all Cummings’ laboratory members. We thank the University of Texas Brackenridge Field Laboratory for animal care facilities.

## Funding

This work was funded by an NSF BEACON Award (26-3509-2650).

## Compliance with ethical standards

This research was conducted without any external financial support.

## Competing interests

The authors have no competing interests.

## Ethical approval

The authors certify that this work followed ethical treatment of animals outlined in their IACUC protocol (AUP-2016-00246).

## Authors’ contributions

Molly Cummings and Kelly Wallace conceived of the study. Kelly Wallace, Mary Ramsey, and Richie Rausch designed and constructed the experimental setup and data collection procedures. Kelly Wallace collected the data and performed statistical analyses. Kelly Wallace and Molly Cummings interpreted the results and wrote the manuscript. All authors gave final approval or publication.

## Notes

https://github.com/kjw2539/Comparative-Cognition-R-scripts

https://github.com/jenkins-cummingslab/ethoStim

