## Supplementary material for "Sex differences in cognitive performance, style and domain relationships in mosquitofish (*Gambusia affinis*)": Electronic Supplementary Materials

*Animal Cognition*

Kelly J. Wallace, Richie T. Rausch, Mary E. Ramsey, Molly E. Cummings

Department of Integrative Biology, University of Texas, Austin, TX, 78712, USA

**Supplementary Figures**

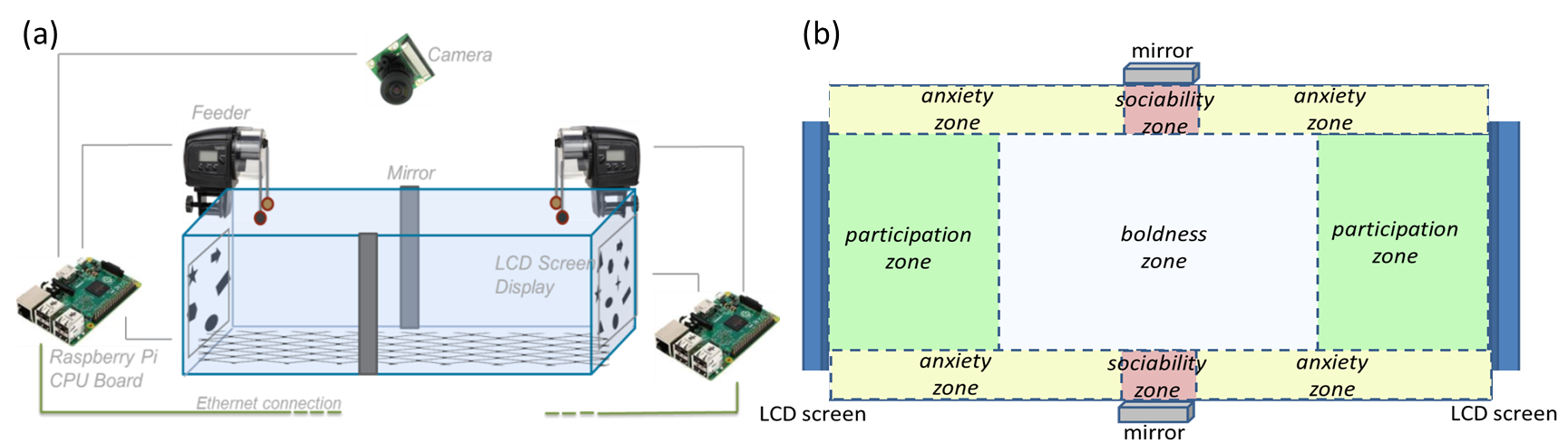

**Supplementary Figure 1**. Apparatus designs for the numerical discrimination assay (a): twelve 40cm x 20cm x 25cm tanks had two Adafruit 7” LCD Displays submerged in mineral oil (to eliminate refraction) at either short end. Above each screen was a modified feeder with a food reward (a marble weight with Cargill Aquaxcel Starter Food® wrapped in netting) and a control stimulus without food. A mirror was attached to the outside of the tank halfway along the long edge to prevent isolation-induced stress and assess sociability tendencies. Meshing placed approximately 1cm above the tank floor allowed for uneaten food to fall to the bottom of the tank and and remain out of reach by the focal fish (to ensure that reward was unavailable when stimuli were absent). Below the meshing was a layer of Deep Blue Professional Ammonia Reducer Pad and a Marineland Bonded Filter Pad for filtration. Sponge filters with air supplies were put in each night and removed by day to maintain healthy conditions. During pilot trials, temperature and dissolved oxygen levels were taken in the morning, afternoon, and evening across multiple days to ensure no changes in condition. Approximate zone designations for assessing cognitive style during the numerical discrimination task (b). A 2cm wide area along long edge of the tank was an area of thigmotaxis (wall-hugging)- proportion of time in this anxiety zone was a measure of anxiety behavior. The participation zone (used to measure associative learning performance) reflects the 10cm closest to the screen (excluding the anxiety zones on either long side). Time spent in the center was used as a measure of boldness, and time spend within a 2cm zone in front of either mirror was used as a measure of sociability.

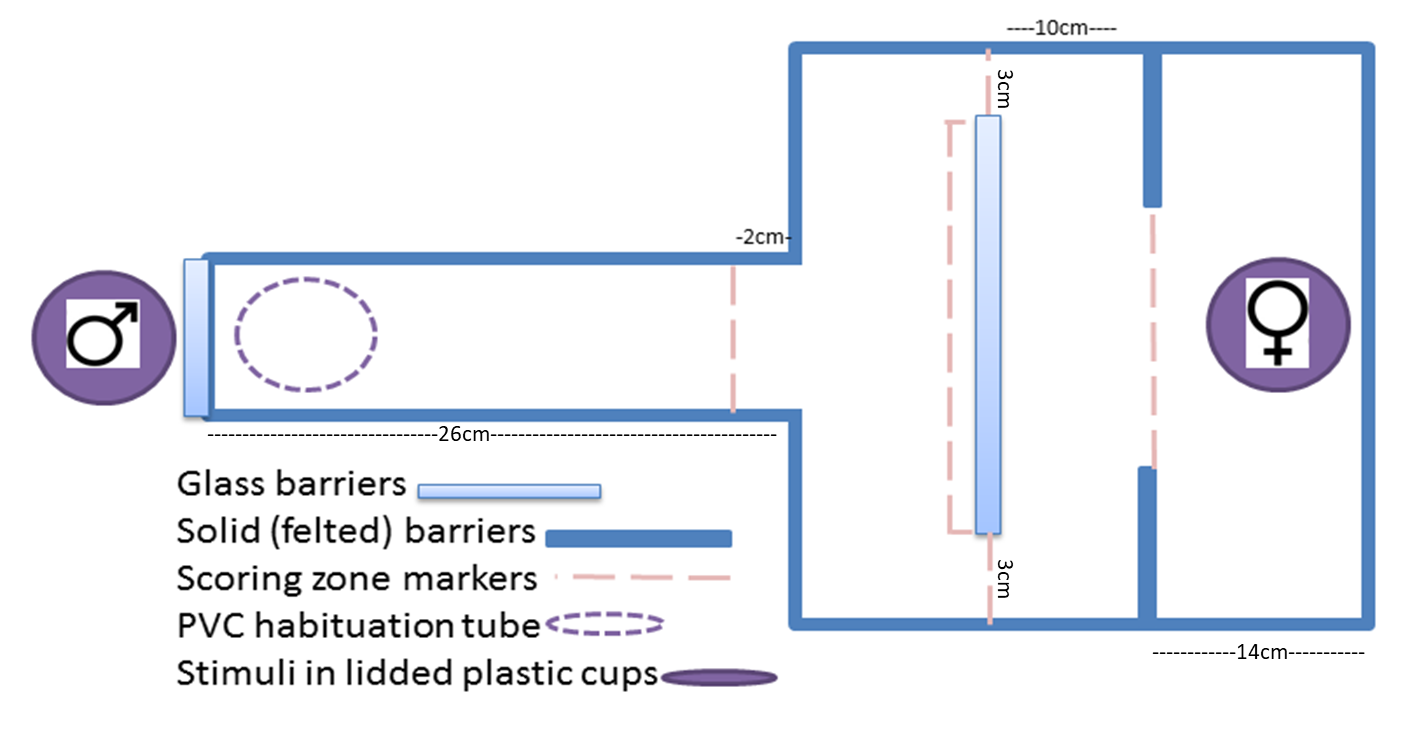

**Supplementary Figure 2.** Apparatus design for the detour maze assay. A male was placed at the near end of the starting alley (26cm x 13.5cm) in a clear lidded plastic cup behind a pane of UV-pass glass. A female was in a clear lidded plastic cup in the reward section (14cm x 31cm). In the middle barrier section (17cm x 31cm) a glass barrier 25 cm wide was placed 10cm in front of the reward zone, centered with ~3cm between the barrier and the long sides of the tank.

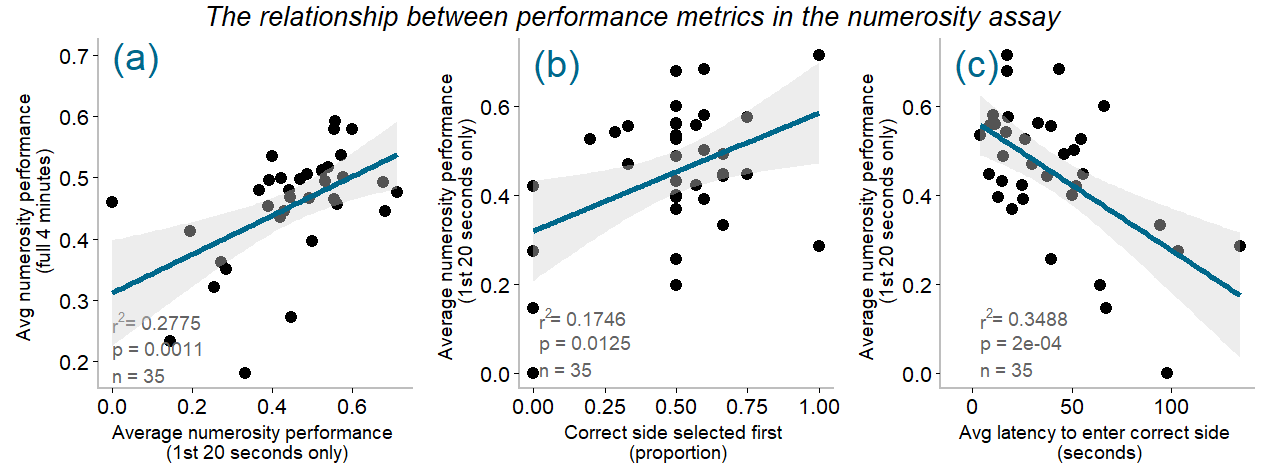

**Supplementary Figure 3.** In the numerical discrimination assay, individual performance was correlated across multiple metrics. Average performance assessed during the 20 second “decision window” significantly represented performance during the entire trial (a), as well as the proportion of trials in which the correct side was the first side chosen/entered (b) and the average latency to enter the correct side (c).

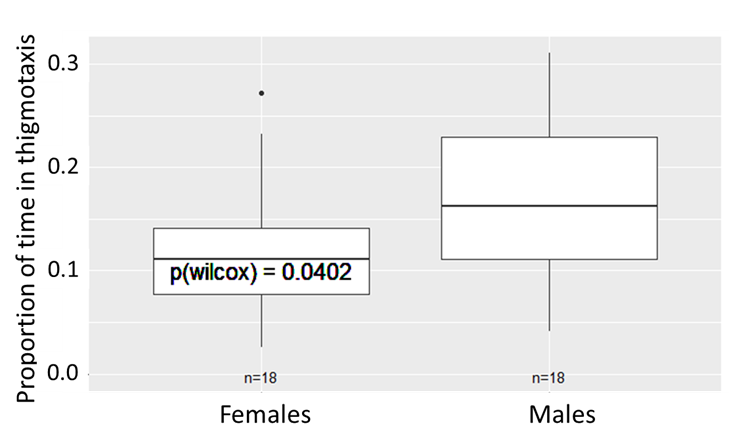

**Supplementary Figure 4.** In the numerical discrimination assay, males show significantly higher average anxiety than females. Anxiety here is defined as the average proportion of the four minute test trials that the fish spent in regions of the tank associated with thigmotaxis, i.e. within 2cm of the long tank edges (see Supplementary Figure 1b).

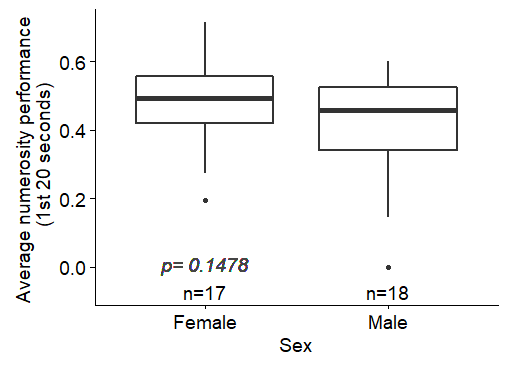

**Supplementary Figure 5.** In the numerical discrimination assay, males and females did not differ in their average performance across 9 testing trials.

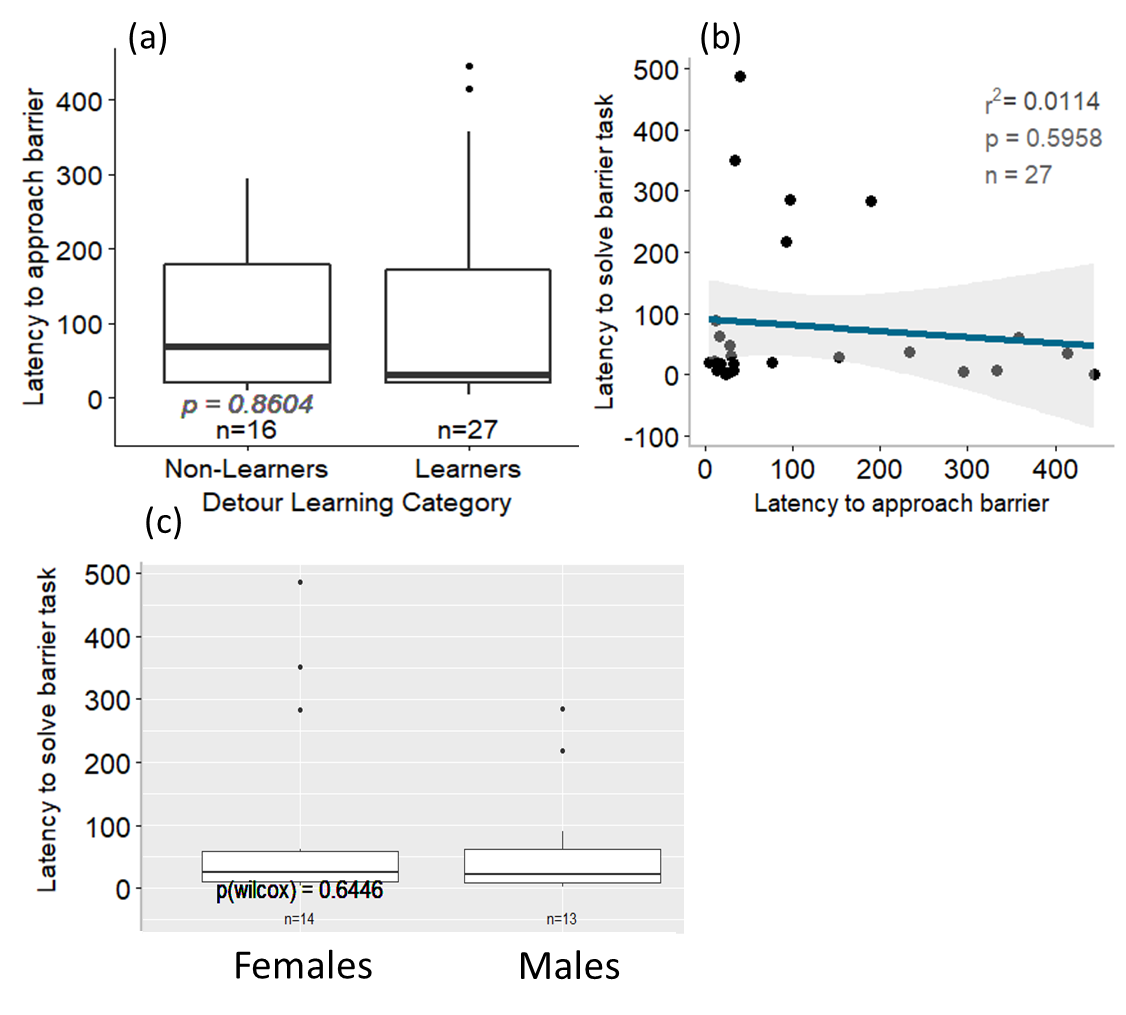

**Supplementary Figure 6***.* Motivation in the detour maze (latency to approach the barrier) does not predict if individuals successfully solve the task (a) or how long it takes for an individual to solve the task (b). No sex difference was seen in latency to solve the detour task (c).

**Supplementary Methods: Experimental Design**

**Methods**

**Housing**

Individuals used in this experiment were wild-caught western mosquitofish *Gambusia affinis* (27 female, 27 male ranging in size from 1.02cm to 3.44cm) from outdoor ponds at Brackenridge Field Laboratories in Austin, TX, housed at the University of Texas at Austin under Institutional Animal Care and Use Committee Protocol AUP-2016-00246 in 35 gallon aquaria at 76-80⁰F(24.4-26.6⁰C) filled with Prime® dechlorinated tap water, and kept on a 13L-11D light cycle. Tank water was constantly circulated and aerated with bubblers. Individuals were fed a mixture of Cargill Aquaxcel Starter Food ® and Tetramin Tropical Flakes ® once daily.

**Cognitive Testing**

Prior to experimental testing, individuals were isolated in 1.25 gallon tanks for 2 days. Each individual then underwent numerical discrimination, detour reaching, and shuttlebox tasks in a randomized balanced order between individuals. All cognition assays were videotaped for subsequent scoring.

**Associative Learning (Numerical Discrimination)**

We employed a numerical discrimination paradigm to assess an individual’s learned association between a quantitative stimulus and a food reward, and additionally we measured several behavioral tendencies: anxiety, sociability, boldness, exploration, reaction time, and activity during the testing trials.

**Numerical Discrimination Tank Design**

Twelve 40cm x 20cm x 25cm 21 liter (16” x 8” x 10” 5.5 gallon) automated tanks were housed at 25.5C. Each tank was surrounded by four fluorescent lights (6am-8pm) suspended 46cm above the shelf and covered with three light filters (Gamcolor 1532, R365, and R4360). The sides of the tank were covered in dark blue felt and two 5cm wide mirrors. The tank floor was covered in a Deep Blue Professional Ammonia Reducer Pad, a Marineland Bonded Filter Pad, and netting placed two cm above the filter pad to prevent uneaten food during the training assays to be unavailable to the fish when training stimuli were absent. Bubblers attached to sponge biofilters circulated and aerated water overnight. To ensure water quality, approximately every 4 days about 500ml (10%) of water was removed and replaced from the tanks. On each of the short ends of the tank a glass well was attached to the outside of the tank containing an Adafruit HDMI 4 Pi: 7” 1024 x 600 LCD Display. The LCD screens presenting the images were submerged in mineral oil wells (14 x 19 cm) attached to the tank wall to eliminate angular-dependent refraction and reflection of the image. The images of shapes presented on the screens (the same stimuli as used by Etheredge et al 2018[1]) controlled for non-numerical quantities: the shapes were positioned to create the same convex hull (overall space the shapes occupy), and all pairs of shapes differed in cumulative surface area by less than 10% (e.g. for an image with more shapes, the size of each individual shape would be slightly smaller). The images were presented on the screen with 2cm distance on either side between the edge of the image and the tank wall, corresponding to the thigmotaxis (“wall-hugging”, a measure of anxiety) region of the tank. Above the screens was a modified Nutrafin Nutramatic 2X fish feeder connected to a Lavolta BPS305 DC Power Supply via an Elegoo L298N DC Motor Driver, holding a food reward (a marble weight plus Cargill Aquaxcel Starter Food wrapped in netting about 20mm in diameter). One Raspberry Pi 5MP Camera Board Module was centered above the tank on average 52cm above the shelf. The camera, feeders, and LCD screens on the tank were connected to two Raspberry Pi 2/3 Motherboard. The four-minute trials were scheduled via Jenkins task manager and ran via a Python script (version 4.8.2). During the trial, the camera would first automatically begin recording, then an image with geometric shapes ranging from five to twelve shapes was presented on the LCD screen. After a 10 second delay, the feeders rotated the food reward below the surface and in front of the “correct” (trained contingency) screen and a control stimulus (a marble the same size as the food reward) in front of the other. The food reward and control stimulus were then were removed after either 230 seconds (habituation) or 10 seconds (training). Version control of the automation Python script available at https://github.com/jenkins-cummingslab/ethoStim for more details.

**Numerical Discrimination Habituation, Training, and Testing**

The morning after initial placement in the numerical discrimination tank, individuals experienced two days of habituation (five trials a day) during which the screens were turned on and displayed a blank (white) image and a food reward was presented for the full trial (four minutes) on both sides. Individuals were then trained for six days (five trials per day at two-hour inter-trial intervals). During these four minute long 2:1 training trials, individuals were presented either 5 versus 10 shapes or 6 versus 12 shapes (black shapes on white background) with the food reward presented in front of the trained contingency for only 10 seconds (while the shapes remained visible for the full 4 minutes). The side (left or right) of image presentation was in a balanced predetermined schedule which did not follow an alternating pattern to avoid training fish to alternating spatial rewards. Half of the individuals were trained to respond to the more numerous quantity, half trained to the less numerous quantity. Following the six days of training, individuals were then tested for three days (three testing trials a day alternated with three additional reinforced trials to prevent extinction effects). The testing stimuli included a novel 2:1 ratio (7 vs 14 shapes) and two additional novel and more challenging ratios a 2:3 of 8 vs 12 shapes, and a 3:4 of 9 vs 12 shapes).

**Numerical Discrimination Scoring**

CowLog 3.0.2 event logging software was used to hand score videos for performance accuracy and behavior. All videos were first gridded to evaluate the time each fish spent in different regions of the test tank (python code can be found at: <https://github.com/kjw2539/make_a_grid.py>.). Human observers scored time spent in each region in front of the stimulus screen (the closest quarter of the tank to the screen, excluding the 2cm on either long side of the tank) as well as proportion of time spent within 2cm of the long tank wall (thigmotaxis measure of anxiety), within 2cm of the 5cm wide mirror (sociability), and in the center of the tank not along the edges (as a measure of boldness). In addition, latency to move to another “zone” of the tank following the image presentation (reaction time), number of unique zones visited during the 4 min test trial (exploration), and total number of transits between zones (activity) were also scored. One author (KW) and twelve undergraduate students independently scored a total of 315 videos. Additionally, scorers scored a subset of videos to assess scorer accuracy: the correlation between eleven scorers was p = 5.41 x 10^-90^ (Single Score Intraclass Correlation), the largest standard deviation in hand scored values measuring proportion time in a zone (left screen, right screen, anxiety, mirror, center) was 1.4% and latency values showed a scorer standard deviation of 0.3 seconds.

**Cognitive Flexibility (Detour maze)**

Cognitive flexibility (“the ability of an individual to change its behavior by developing new responses to novel stimuli or altering existing responses to familiar stimuli[2]”) has frequently been tested in varied taxa such as chickens, quail, dogs, dingoes, and quokkas using a transparent detour reaching task[3]. In a typical detour reaching task, individuals must navigate around a transparent barrier to reach a reward. Here, we employ a socially-motivated single-trial transparent detour maze.

**Cognitive Flexibility Tank Design**

The detour maze experimental apparatus was a tank filled with 13cm of water. The lighting environment consisted of two fluorescent lights, hung on either long side of the tank at approximately 22cm high from the base and covered with the following light filters: Diffuse R102, R4360, Lee201, and either Lee298 or Gamcolor 1532. Lux measurements read between 20 (reward zone) and 95 (end of alley), with an average lux of 55 (reader settings at Auto 2000). Because research has recorded that female *Gambusia affinis* shoal with other females in the presence of males to reduce male harassment [4,5] we use a male as a social activator and a female as a social reward. The male “social activator” was placed at the near end of the teal-felted starting alley section (26cm x 13.5cm) in a clear lidded plastic cup behind a pane of UV-pass glass. At the far end in the green-felted reward zone (14cm x 31cm) a female “social reward” was in a clear lidded plastic cup. In the blue-felted barrier zone (17cm x 31cm) a glass barrier 25 cm wide was placed 10cm in front of the reward zone, centered with ~3cm between the barrier and the long sides of the tank. At the start of the trial, the focal individual was placed in starting alley near the male social activator in a PVC tube (10 cm in diameter) for a five-minute habituation, then the PVC pipe was removed and the individual was allowed to swim freely for 10 minutes before being returned to their home tank. Videos of the trial were recorded using Debut Video Capture Software and LifeCam cameras.

**Cognitive Flexibility Scoring**

Videos were first renamed, removing the identity of the focal individual to prevent scoring bias. Scorers recorded the time at which an individual left the starting alley, approached the barrier, navigated around the barrier, and entered the reward zone. Subsequently to the focal fish first entering the reward zone, scorers recorded the amount of time spent in the reward zone. Detour videos were independently scored by hand by two undergraduate student scorers and compared to scores of the co-author (KW) scorer. Correlation between the two student scorers and the co-author scorer were p = 4.83 10^-45^ and p = 2.03 x 10^-35^. Average difference in scorer latency when validated was 4.17 seconds.

**Temporal learning (Shuttlebox)**

While the shuttlebox paradigm has been extensively used in animal psychology studies [6], a socially motivated shuttlebox assay in fish only been recently employed [7]. In this socially motivated shuttlebox assay, individuals are presented a video of a conspecific shoal group at alternating ends of the experimental tank to assess temporal learning. The shuttlebox experimental apparatus consisted of a 52 x 26 tank filled to 10cm of water. On either long side a fluorescent light covered in light filters: a diffusion filter, a Gamcolor 1532, R365, and R4360 were placed. The sides of the tank were covered in tan felt and the bottom in light blue felt to prevent reflection.

During a 5-minute habituation period an individual was placed in the tank and allowed to swim freely while two Adafruit HDMI 4 Pi: 7” 1024 x 600 LCD Displays showed a video of an empty experimental tank apparatus covered in tan felt. Following the habituation, one screen (either the left screen or right screen, balanced across individuals) would present a stimulus presentation: a 20 second video recording of 5 conspecific female *Gambusia affinis* in the same experimental tank as was played during the habituation. During this time, the opposite screen continued to show the empty tank. After 20 seconds, both screens showed the empty tank video for a 90 second inter-stimulus interval. Following the inter-stimulus interval, the conspecific video would play on the opposite side as previously played. These alternating stimulus presentations separated by 90 second inter-stimulus intervals continued for a period of one hour, at which time the fish was removed from the apparatus and placed back in its home tank. Videos of the trial were recorded using Debut Video Capture Software and LifeCam cameras.

**Supplementary Table 1. Shuttlebox Experimental Design*.***  In the shuttlebox assay, a video of a 5-female conspecific shoal is alternated between screens at opposite ends of a tank followed by an inter-stimulus-interval. The screen opposite the shoal video plays a video of an empty tank.

|  | *Habituation*  *(5 min)* | *Stimulus Presentation (20 s)* | *ISI*  *(90 s)* | *Stimulus Presentation (20 s)* | *ISI*  *(90 s)* | *Stimulus*  *Presentation (20 s)* |
| --- | --- | --- | --- | --- | --- | --- |
| *Left Screen*  *video* | Empty | Empty | Empty | **Conspecific** | Empty | Empty |
| *Right Screen*  *video* | Empty | **Conspecific** | Empty | Empty | Empty | **Conspecific** |

**Shuttlebox Scoring:** The shuttlebox videos were hand scored using a python program developed in lab (see https://github.com/lukereding/shuttlebox/blob/master/track/shuttlebox_hand_track.py, developed by Luke Reding February 2018). The program pulled a frame from the video every 5 seconds and displayed it to a student scorer, who located and clicked the position of the center of the fish on the image.

### **Supplementary Methods: Numerical Discrimination Setup**

The list of materials and components needed to support running a round include:

| Component | Description | Quantity |
| --- | --- | --- |
| Fish Tank | Standard 40x20x25cm tank (housed at 25.5⁰C and filled with Prime® dechlorinated tap water) modified with custom built plexiglass container on each end to hold a screen submerged in mineral oil. Each tank was oxygenated from 5pm-9am. | 6 |
| Raspberry Pi (RPi) 3 Model B v1.2 | Small and affordable computer that provides interfaces for connecting screen (HDMI), USB devices, PiCamera, and series of GPIO pins that can drive various voltages to control other electronics. | 12 |
| LaVolta BPS-305 | Variable 30V 5A DC power supply | 1 |
| L298N | Dual H-Bridge motor driver which allows speed and direction control of two DC motors at the same time. | 6 |
| Raspberry Pi Camera Board v1.3 | Camera that connects directly to the RPi’s camera serial interface connector. 5MP camera module capable of 2592 x 1944 pixel static images, and also supports 1080p @ 30fps, 720p @ 60fps and 640x480p 60/90 video. | 6 |
| Nutrafin Nutramatic 2X Fish Feeder | Fish feeder with modifications. These feeders are typically battery powered but were modified to expose the power and ground wires connected to the motor which then could be driven by an external power supply. The feed delivery system was modified to add cables to power and ground via the L288N Motor Driver | 12 |
| Adafruit 2407 | 7” Mini HDMI monitor with built-in touchscreen. | 12 |
| TP-Link TL-SG105 | 5-port gigabit desktop switch^[1]^ | 3 |
| Seagate SRD00F1 | 4TB portable hard drive connected to alien for networked video file storage. | 1 |
| alien (master node) | This is a desktop PC running Ubuntu that from a firewall perspective is able to communicate with all 24 RPis, it also hosts the master instance of the Jenkins software including the webpage interface. | 1 |
| Rack | 3 level rack that can accommodate 2 tanks plus equipment per level | 1 |
| GE 18-Inch Basic Fluorescent Light Fixture 16466 | Each tank was surrounded by four fluorescent lights (6am-8pm) suspended 46cm above the shelf and covered with three light filters (Gamcolor 1532, R365, and R4360). Diffuse light from other shelves/ceiling/etc. was reduced via black fabric curtains. |  |

### Physical Setup

### Each round of trials consists of up to 6 fish with 1 fish per tank. Each tank requires 2 fish feeders, 2 screens, 2 RPis, 1 PiCamera, and 1 motor driver chip. All the RPIs are networked together using a series of Ethernet switches connected to our master PC which provides 4TB of network storage via USB connected portable hard drive Using firewall settings, only the master PC can access the RPis over the network (i.e. SSH). A diagram of a single setup if shown in the figure below: WE WILL NEED A FIGURE TITLE & Legend for this.

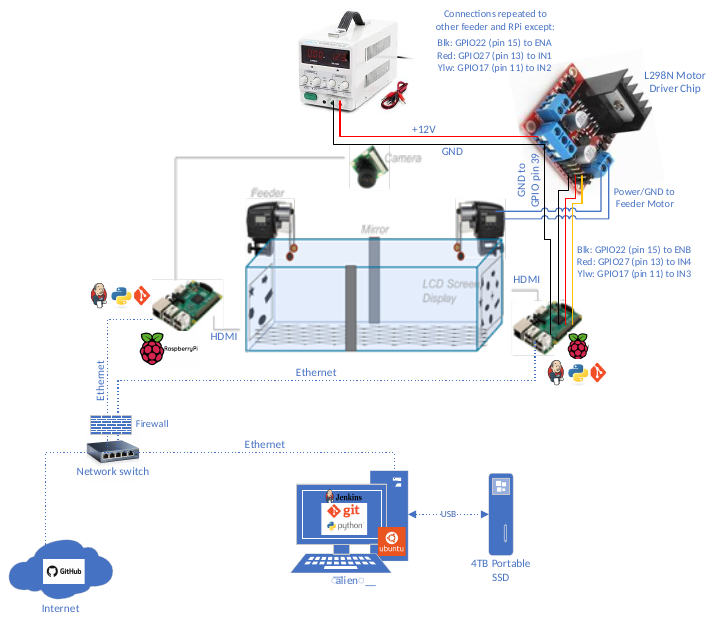

### Scripts

All of the scripts are available on GitHub at <https://github.com/jenkins-cummingslab/ethoStim>. The scripts are primarily written in Python and configuration files are in the JSON format. The Python scripts are all written to execute with Python 2.7 (i.e. no guarantees with Python3). The scripts were developed to be OS-agnostic. Development and testing were primarily performed in a Windows 10 environment, but ultimately executed on either an Ubuntu or Raspian (both Linux-based distros) environment. Python can be downloaded and installation instructions found at <https://www.python.org/downloads/>. The following are Python modules that may not be included with a standard Python installation:

| Module | Use | Download Location |
| --- | --- | --- |
| pygame | Displaying image to screens | <https://pypi.org/project/Pygame/> |
| picamera | Interfacing with the PiCamera device | <https://pypi.org/project/picamera/> |
| RPi.GPIO | Drive GPIO pins on RPi that are used to control motors in the fish feeders | <https://pypi.org/project/RPi.GPIO/> |
| netifaces | Used to get addresses of network interfaces | <https://pypi.org/project/netifaces/> |

Many of the scripts utilize or modify configuration files. All of the configuration files are in the JSON format (<https://www.json.org/>). This format was chosen because it’s easy for humans to read and write and easy for machines to parse and generate. For just about any modern programming language, there are high-quality libraries available to support JSON reading and writing.

The majority of these scripts are rarely, if ever, run by a human in our setup. There is a Jenkins job that wraps this behavior that is either triggered automatically from another Jenkins job or the user is provided a simple webpage form for the user to fill out.

NOTE: The repository is a fork from <https://github.com/ietheredge/ethoStim>. The original repository was used in early versions of this experiment, but has evolved and been optimized since then. Refer to the repository commit history for details on changes since the fork.

The following sections provide an overview of the key scripts.

#### trial.py

This script is run on a RPi and provides support for controlling a fish feeder, displaying images on a screen, and interfacing with a PiCamera. Based on configuration parameters, the script utilizes a series of timers to perform the desired functions at the desired trigger time and for the desired duration. The fish feeder is controlled by utilizing GPIO pins available on the RPi to driver the motor directly. It supports driving the motor in either the clockwise or counter-clockwise directions and also can support variable speed via a pulse-width modulation technique. The configuration parameters are either presented as input arguments or read through a specifically named JSON configuration file. The command line usage is shown below:

```
>>python trial.py --help
usage: trial.py [-h] [-a CWTIME] [-b CCWTIME] [-f FISH] [-ts THATPISTIMULUS]
                [-ps PISTIMULUS] [-cs CORRECTSIDE] [-d DAY] [-s SESSION]
                [-fs FEDSIDE] [-x SEX] [-p PROPORTION] [-sp SPECIES]
                [-sl FISHSTANDARDLENGTH] [-r ROUND] [-fd] [-c] [-m STARTTIME]
                [-sd STARTDELAY] [-sync]
optional arguments:
  -h, --help            show this help message and exit
  -a CWTIME, --cwtime CWTIME
                        Time is seconds that motor(s) is on in clockwise dir
  -b CCWTIME, --ccwtime CCWTIME
                        Time is seconds that motor(s) is on in counter
                        clockwise dir
  -f FISH, --fish FISH  ID of fish in tank
  -ts THATPISTIMULUS, --thatpistimulus THATPISTIMULUS
                        numerosity stimulus being shown on the other raspberry
                        pi in the tank
  -ps PISTIMULUS, --pistimulus PISTIMULUS
                        stimulus being presented with this raspberry pi
  -cs CORRECTSIDE, --correctside CORRECTSIDE
                        stimulus side on which the correct stimulus is being
                        presented
  -d DAY, --day DAY     experiment day, e.g. 1-7
  -s SESSION, --session SESSION
                        trial session, e.g. 1-4
  -fs FEDSIDE, --fedside FEDSIDE
                        side feed on/conditioned side
  -x SEX, --sex SEX     fish sex
  -p PROPORTION, --proportion PROPORTION
                        ratio that is being presented this trial
  -sp SPECIES, --species SPECIES
                        species name
  -sl FISHSTANDARDLENGTH, --fishstandardlength FISHSTANDARDLENGTH
                        standard length of the
  -r ROUND, --round ROUND
                        training round
  -fd, --feed           feed with this stimulus
  -c, --camera          do you want to record using this pi?
  -m STARTTIME, --startTime STARTTIME
                        time since epoch that you want to start your trial
  -sd STARTDELAY, --startDelay STARTDELAY
                        Number of seconds to delay start (for jenkins this
                        should be 180), NA if sync is false
  -sync, --dosync       do you want to attempt to time sync using startTime
                        and startDelay?
>>
```

The contents of an example trial_config.json is shown below:

```
{
  "SLEEP_AFTER_CAMERA_START_SECS": "30",
  "FEED_DURATION_SECS": "10",
  "FEED_DURATION_SECS_HABITUATION": "1800",
  "FEED_DELAY_SECS": "10",
  "TRIAL_DURATION_SECS": "275",
  "TRIAL_DURATION_SECS_HABITUATION": "1855",
  "VID1_LEN_SECS": "275",
  "VID2_LEN_SECS": "30",
  "INCLUDE_VID2": "0",
  "START_MAX_MINS_IN_PAST": "60"
}
```

At the conclusion of a successful execution of the script, an H264 encoded video file is generated with the following naming convention (uses many of the command line arguments):

species_round_sl_sex_fishid_day_session_stim_thatpistimulus_proportion_fedside_correctside.h264

An example would be:

gambusia_22_216_male_Garrison_1_2_0_0_0_both_B.h264

NOTE: Although most video players will support playback of an H264 encoded video file, many will lack full functionality (pause, rewind, fast-forward, etc.). Tools like ffmpeg (<https://www.ffmpeg.org/>) can be used to “wrap” an H264 video file into container such as MP4.

#### turn_screen_on.py.

This script can be used to simply turn on the screen and display one of the standard images provided in the ethoStim repository (e.g. 0.png, 5.png, etc.).

```
>>python turn_screen_on.py --help
usage: turn_screen_on.py [-h] [-l LEN] [-i IMAGE]
optional arguments:
  -h, --help            show this help message and exit
  -l LEN, --len LEN     How long do you want the screen on (secs)
  -i IMAGE, --image IMAGE
                        Image to show on screen
>>
```

#### add_to_fish_json.py

This is a helper script used to add a fish and details about the fish that Jenkins will use to generate and schedule jobs to run a trial. It interacts with fish.json which is available in the repository. The JSON file could be modified by hand, but this script will ensure JSON formatting is maintained.

```
>>python add_to_fish_json.py --help
usage: add_to_fish_json.py [-h] -f FISH -sp SPECIES -s SEX -sl
                           FISHSTANDARDLENGTH -n NODE -c CAMNODE
optional arguments:
  -h, --help            show this help message and exit
  -f FISH, --fish FISH  Name of fish.
  -sp SPECIES, --species SPECIES
                        Species of fish.
  -s SEX, --sex SEX     Sex of fish
  -sl FISHSTANDARDLENGTH, --fishstandardlength FISHSTANDARDLENGTH
                        Standard length of fish
  -n NODE, --node NODE  Node without camera
  -c CAMNODE, --camnode CAMNODE
                        Node with camera
>>
```

The contents of an example fish.json could be:

```
{
    "Alexandra": {
        "cam_node": "pluto",
        "fishstandardlength": "251",
        "node": "ceres",
        "sex": "female",
        "species": "nigrensis"
    },
    "Alyssa": {
        "cam_node": "uranus",
        "fishstandardlength": "386",
        "node": "neptune",
        "sex": "female",
        "species": "nigrensis"
    },
    "Amanda": {
        "cam_node": "haumea",
        "fishstandardlength": "301",
        "node": "makemake",
        "sex": "female",
        "species": "nigrensis"
    },
}
```

#### del_fish_from_json.py

This is a helper script used to delete a fish from fish.json. The JSON file could be modified by hand, but this script will ensure JSON formatting is maintained. This is primarily used for cleanup purposes and maintaining only valid fish, but it also is important because it can be used to prevent Jenkins from creating jobs for an invalid fish (ie someone typos an old fish name for round configuration). If a fish is not in fish.json, Jenkins will error gracefully and notify the user.

```
>>python del_fish_from_json.py --help
usage: del_fish_from_json.py [-h] -f FISH
optional arguments:
  -h, --help            show this help message and exit
  -f FISH, --fish FISH  Name of fish.
>>
```

#### add_to_pies_json.py

This is a helper script used to add a RPi and details about the RPi that Jenkins will use to generate and schedule jobs to run a trial. It interacts with pies.json which is available in the repository. The JSON file could be modified by hand, but this script will ensure JSON formatting is maintained.

```
>>python add_to_pies_json.py --help
usage: add_to_pies_json.py [-h] -p PI -cw CWTIME -ccw CCWTIME -cd COPYDELAY
optional arguments:
  -h, --help            show this help message and exit
  -p PI, --pi PI        Name of pi.
  -cw CWTIME, --cwtime CWTIME
                        Clockwise time (secs).
  -ccw CCWTIME, --ccwtime CCWTIME
                        Counter clockwise time (secs)
  -cd COPYDELAY, --copydelay COPYDELAY
                        Delay before copy to shared drive (secs)
>>
```

The contents of an example pies.json could be:

```
{
    "haumea": {
        "ccwtime": "0.8",
        "copydelay": "1980",
        "cwtime": "0.8"
    },
    "pluto": {
        "ccwtime": "3.8",
        "copydelay": "1800",
        "cwtime": "3.8"
    },
    "uranus": {
        "ccwtime": "3.8",
        "copydelay": "1620",
        "cwtime": "3.8"
    }
}
```

#### del_pi_from_json.py

This is a helper script used to delete a RPi from pies.json. The JSON file could be modified by hand, but this script will ensure JSON formatting is maintained. This is primarily used for cleanup purposes and maintaining only valid RPis, but it also is important because it can be used to prevent Jenkins from creating jobs for an invalid RPi (ie someone typos an old RPi name for round configuration). If a RPi is not in pies.json, Jenkins will error gracefully and notify the user.

```
>>python del_fish_from_json.py --help
usage: del_fish_from_json.py [-h] -f FISH

optional arguments:
  -h, --help            show this help message and exit
  -f FISH, --fish FISH  Name of fish.
>>
```

#### mod_top_json.py

This is a helper script used to define the top level parameters for a round trials. The parameters are stored in a JSON file named top.json. The script will overwrite an existing top.json with the new parameters. The JSON file could be modified by hand, but this script will ensure JSON formatting is maintained.

```
>>python mod_top_json.py --help
usage: mod_top_json.py [-h] -sd STARTDATE -r ROUND -hs HSCHED -ls LSCHED -h1
                       HFISH1 -h2 HFISH2 -h3 HFISH3 -l1 LFISH1 -l2 LFISH2 -l3
                       LFISH3
optional arguments:
  -h, --help            show this help message and exit
  -sd STARTDATE, --startdate STARTDATE
                        Date to start scheduling trials, format is MM/DD.
  -r ROUND, --round ROUND
                        A number.
  -hs HSCHED, --hsched HSCHED
                        Which high schedule to use (e.g. H1, H2, H3)
  -ls LSCHED, --lsched LSCHED
                        Which low schedule to use (e.g. H1, H2, H3)
  -h1 HFISH1, --hfish1 HFISH1
                        1st Fish that will be assigned H schedule
  -h2 HFISH2, --hfish2 HFISH2
                        2nd Fish that will be assigned H schedule
  -h3 HFISH3, --hfish3 HFISH3
                        3rd Fish that will be assigned H schedule
  -l1 LFISH1, --lfish1 LFISH1
                        1st Fish that will be assigned L schedule
  -l2 LFISH2, --lfish2 LFISH2
                        2nd Fish that will be assigned L schedule
  -l3 LFISH3, --lfish3 LFISH3
                        3rd Fish that will be assigned L schedule
>>
```

The contents of an example top.json could be:

```
{
    "h_schedule": "H1",
    "l_schedule": "L1",
    "mapping": {
        "H": {
            "fish1": "amber",
            "fish2": "andrea",
            "fish3": "amelia"
        },
        "L": {
            "fish1": "becca",
            "fish2": "bella",
            "fish3": "beth"
        }
    },
    "round": "1",
    "startDate": "07/15"
}
```

#### gen_trial_jobs.dsl

This is the only non-Python script provided in the repository. This is a Groovy script. The Groovy programming or scripting language is very Python-like and can also interact directly with Java code and libraries.

This script is used by Jenkins to generate a round of trials. This job will NOT run outside of the Jenkins environment, it’s only purpose is to generate Jenkins jobs. It reads in and parses a series of JSON configuration files that describe the round, fish, RPi, and schedule parameters and generates a wrapper job for each individual trial that calls a series of parameterized jobs that actually execute the trial (i.e. runs trial.py) on a specific RPi, copies the video file to a network share, and converts the resulting encoded H264 video file to MP4 video file. The following is a list of all the JSON files the script requires:

top.json

fish.json

pies.json

H1.json

H2.json

H3.json

L1.json

L2.json

L3.json

NOTE: H#.json and L#.json define the schedule parameters. H is for fish being trained to the high schedule and L is for fish being trained to low schedule. We created 3 sets of high schedules and 3 sets of low schedules. A high schedule provides a food reward on the side of the tank that is displaying image with more shapes during training. A low schedule provides a food reward on the side of the tank that is displaying image with fewer shapes during training.

### Jenkins

Jenkins is an open-source server-based automation framework that supports continuous integration and delivery in the software development process. It has a large and growing community providing online support and developing plug-ins to extend the base functionality. Users include Raytheon, Qualcomm, Facebook, Dell, LinkedIn, Netflix, Google, Carfax, and many more. The basic (and most common) tenet of Jenkins is to monitor changes to a source code repository to trigger build verification, testing, and artifact deployment “jobs” along with redundant verification (e.g. nightly build and test) and additional testing needs (e.g. adversarial, stress, long-duration testing). More generally, if a task can be automated, Jenkins can be used and is a solid choice for executing and managing automation needs.

In past iterations of the numerosity setup, a human would access each RPi and deploy a Cron job (i.e. time-based scheduler) to schedule trials. Throughout a round, a human had to routinely access the RPi to move video files off the RPi due to limited memory. Keeping track if a job had run successfully and overall health of the experiment was difficult at best. The overall process from job scheduling to video file management was error prone, time consuming, and required ample human interaction and coordination. Jenkins allows us to fully automate this process and effectively remove the human error component which in turn allowed the human to focus on other aspects of the research. Using Jenkins, we were able to easily network all of the RPis together and manage them from a central “master” node. Utilizing Jenkins artifact deployment features, we were able to centralize storage off the RPis. From a single webpage hosted by the master node, a user can configure and schedule a round for all the RPis, monitor the health of the setup, monitor the status of jobs, and access video and log files. The webpage user interface is easy to use and monitor and for most cases, simply involves filling out a pre-formatted form and pressing a button.

Jenkins user documentation and download can be found at <https://jenkins.io/download/>. For our setup, we are using Jenkins version 2.46.3 hosted on our master node desktop computer running Ubuntu LTS 14.04.5 LTS. The following table provides a non-inclusive list of plug-ins installed focusing primarily on plug-ins that are not typically available with a base installation.

| Plug-In | Version | Short Description |
| --- | --- | --- |
| Artifact Deployer | 0.33 | Makes it possible to deploy artifacts from workspace to output directories |
| Build Timeout | 1.18 | Allows builds to be automatically terminated after the specified amount of time has expired |
| Build Timestamp | 1.0.1 | Adds BUILD_TIMESTAMP to Jenkins variables and system properties |
| Credentials Binding | 1.10 | Allows credentials to be bound to environment variables for use from the miscellaneous build steps |
| description setter plugin | 1.10 | Sets the description for each build, based upon a RegEx test of the build log files |
| Email Extension | 2.52 | Replacement for Jenkins’s email publisher |
| Environment Injector | 2.1 | Makes it possible to set an environment for the builds |
| Git | 3.0.1 | Integrates Git with Jenkins |
| GitHub | 1.23.1 | Integrates GitHub to Jenkins |
| Gradle | 1.25 | Allows Jenkins to invoke Gradle build scripts directly |
| Groovy | 1.30 | Allows Jenkins to invoke Groovy build scripts directly |
| Job DSL | 1.53 | Allows Jobs and Views to be defined via DSLs |
| JobDelete Builder Plugin | 1.0 | Delete job in build step |
| Maintenance Jobs Scheduler Plugin | 0.1.0 | Explains how to write a Jenkins plugin |
| Node and Label parameter plugin | 1.7.2 | Allows to dynamically select the node on which a job should be executed |
| Persistent Parameter Plugin | 1.1 | String, text, Boolean, and choice parameters with default values set from the previous build (if any) |
| Pipeline | 2.4 | A suite of plugins that lets you orchestrate automation, simple or complex |
| Purge Build Queue Plugin | 1.0 | Provides an easy mechanism to purge the build queue |
| Purge Job History Plugin | 1.1 | Provides the ability to purge all the build records of a job via a CLI command or via the UI |
| Rebuilder | 1.25 | Used for rebuilding a job using the same parameters |
| SSH Slaves | 1.11 | Allows to launch agents over SSH, using a Java implementation of the SSH protocol |
| Timestamper | 1.8.7 | Adds timestamps to the Console Output |
| Workspace Cleanup | 0.32 | Deletes the project workspace after a build is finished |

To provide some context to how Jenkins actually works, a sample job is created below with screenshots. The example job includes running a Python script (i.e. hello.py) that is pulled down from GitHub that needs to execute on a specific computer (i.e. node) everyday at 6am. The script produces an output file (i.e. out.log) that needs to be stored for future analysis.

1. Go to Jenkins homepage, accessed through web browser such as FireFox, Internet Exporer, or Chrome.

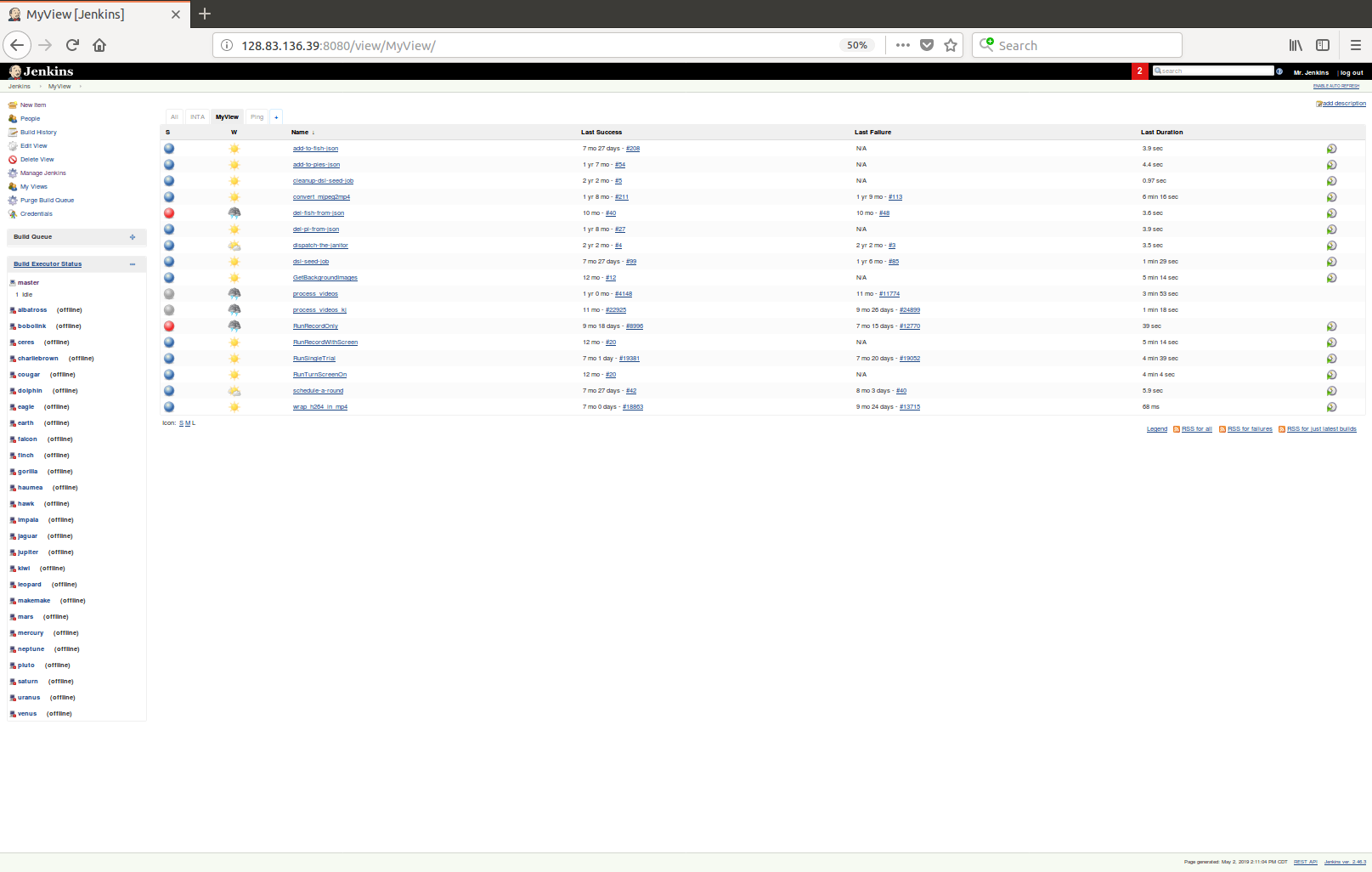

homepage

actions

available nodes

jobs

2. Click on New Item

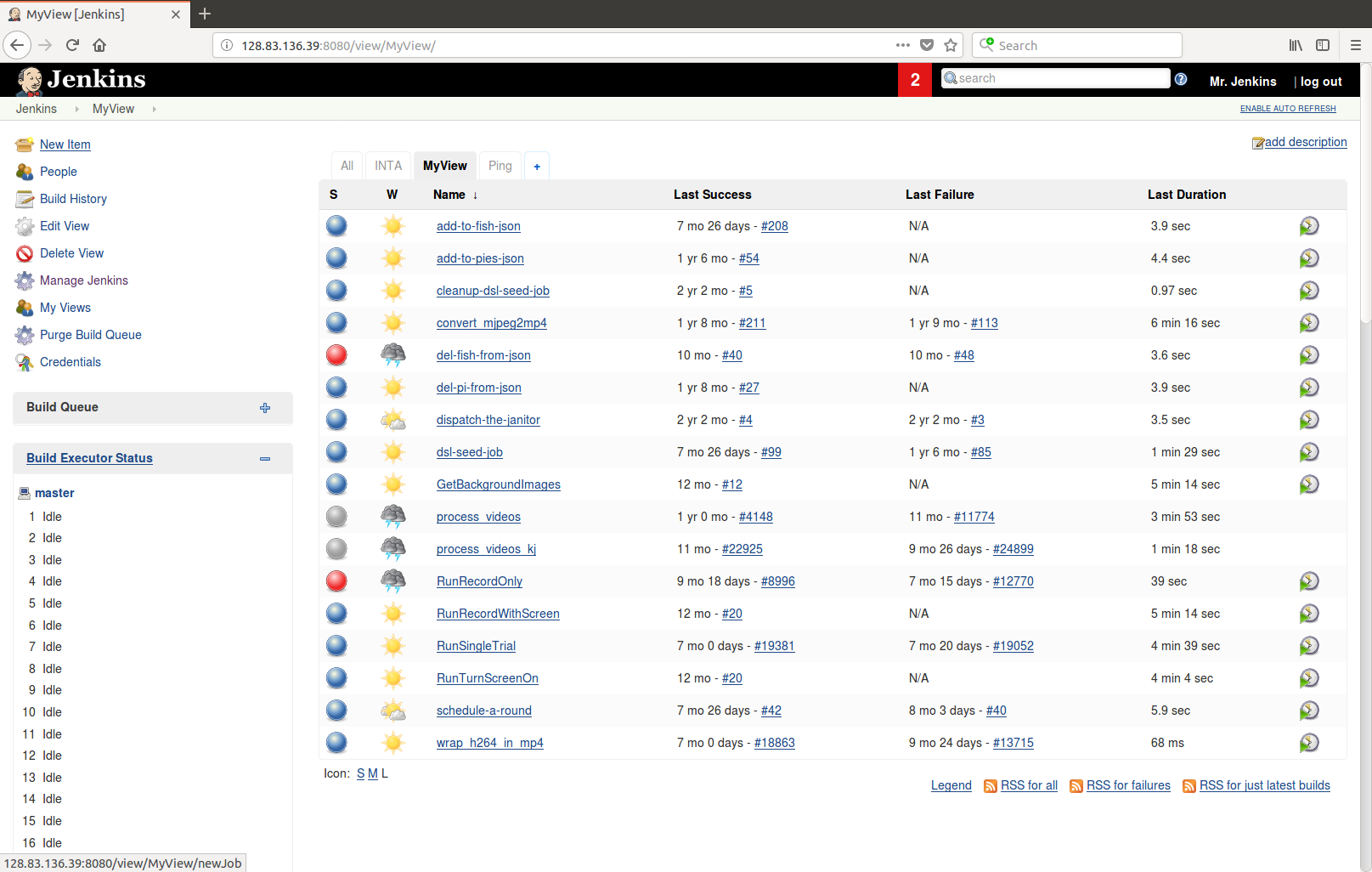

3. Enter a name for the job, type of job, and click OK.

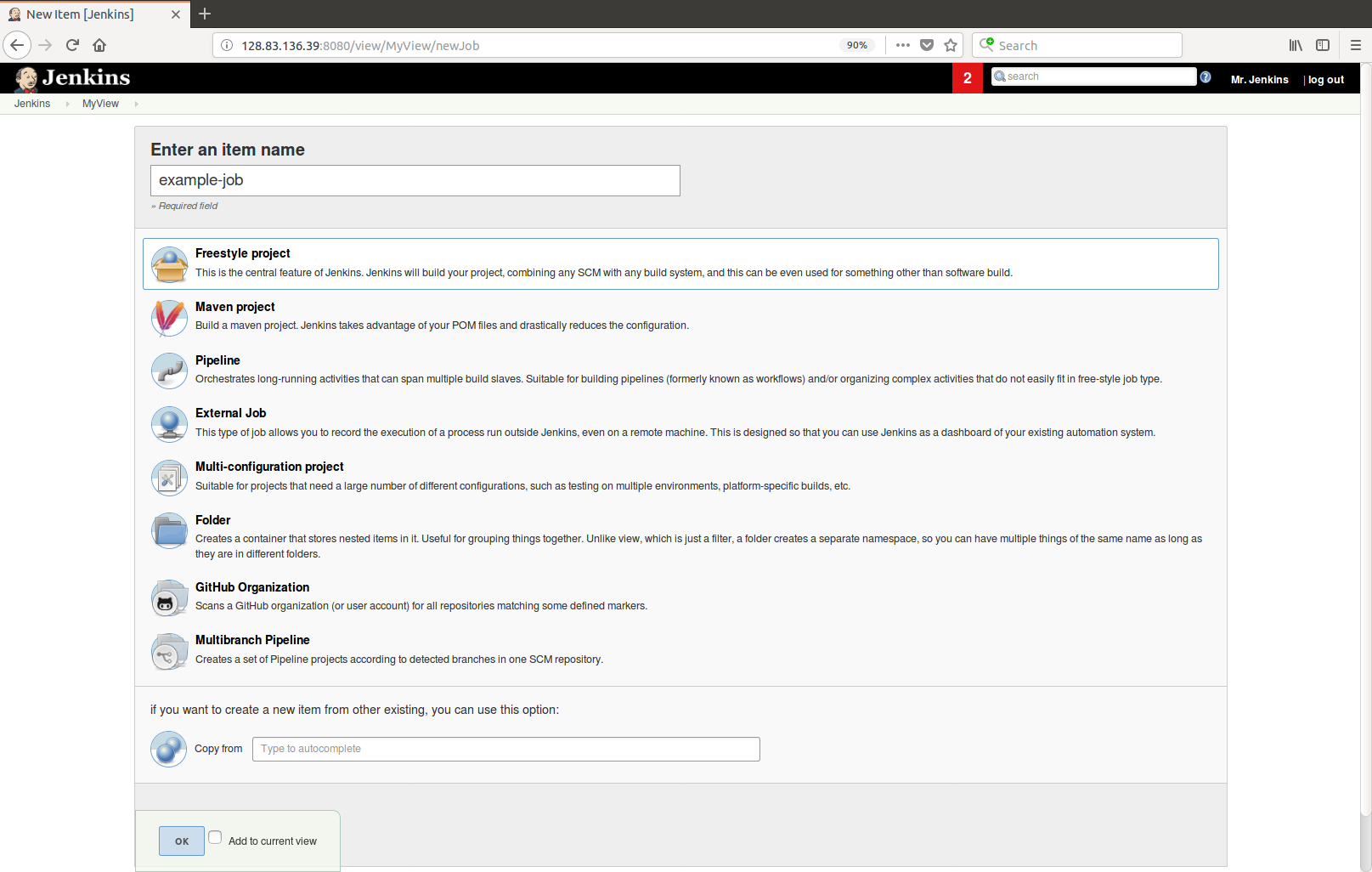

4. Configure the General settings of the job including job description, number of builds to keep and/or how long to keep builds, any job parameters, the node (i.e. computer) to run the job on, etc.

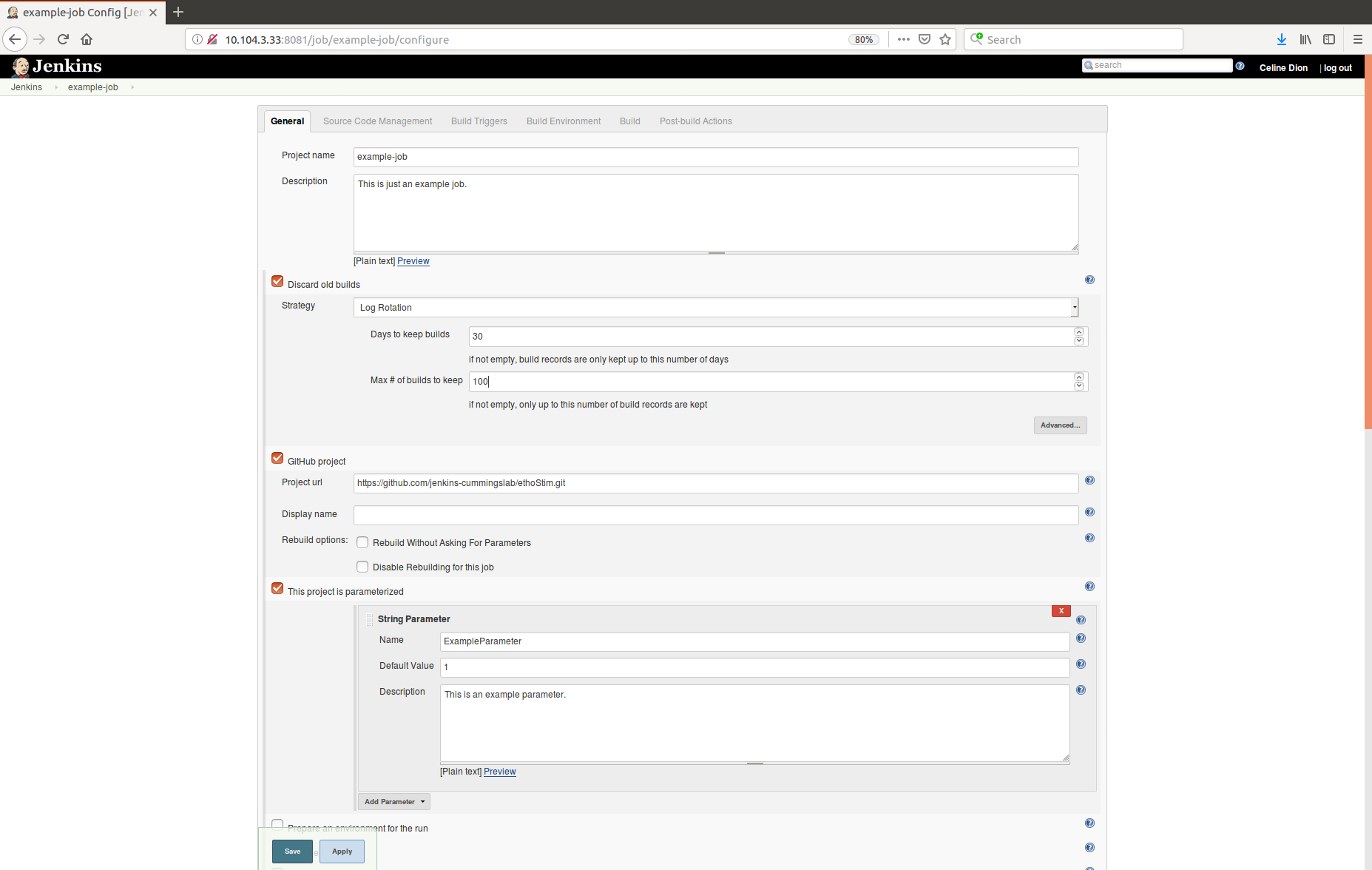

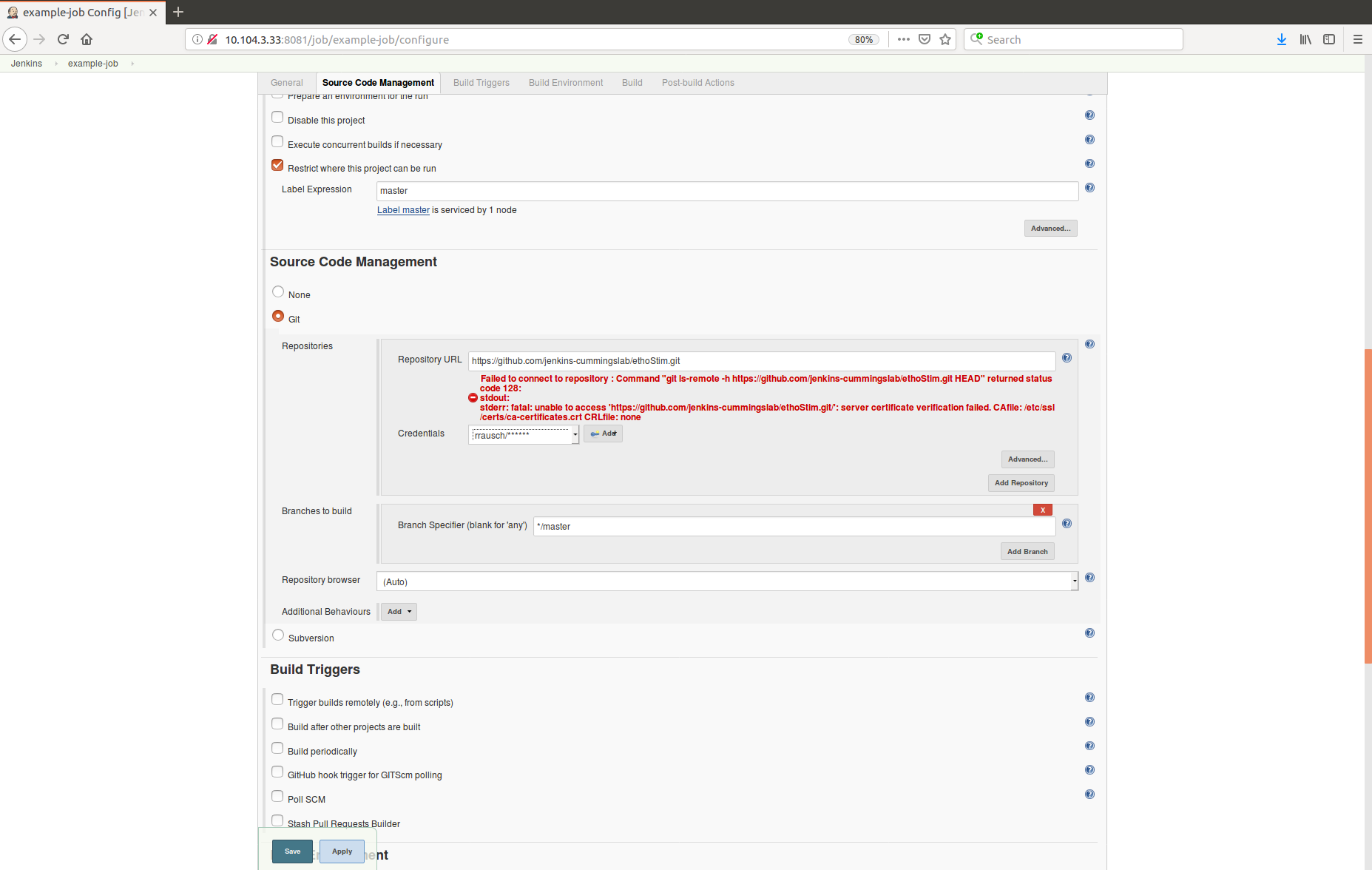

5. Configure the Source Code Management settings such as the GitHub URL and associated credentials.

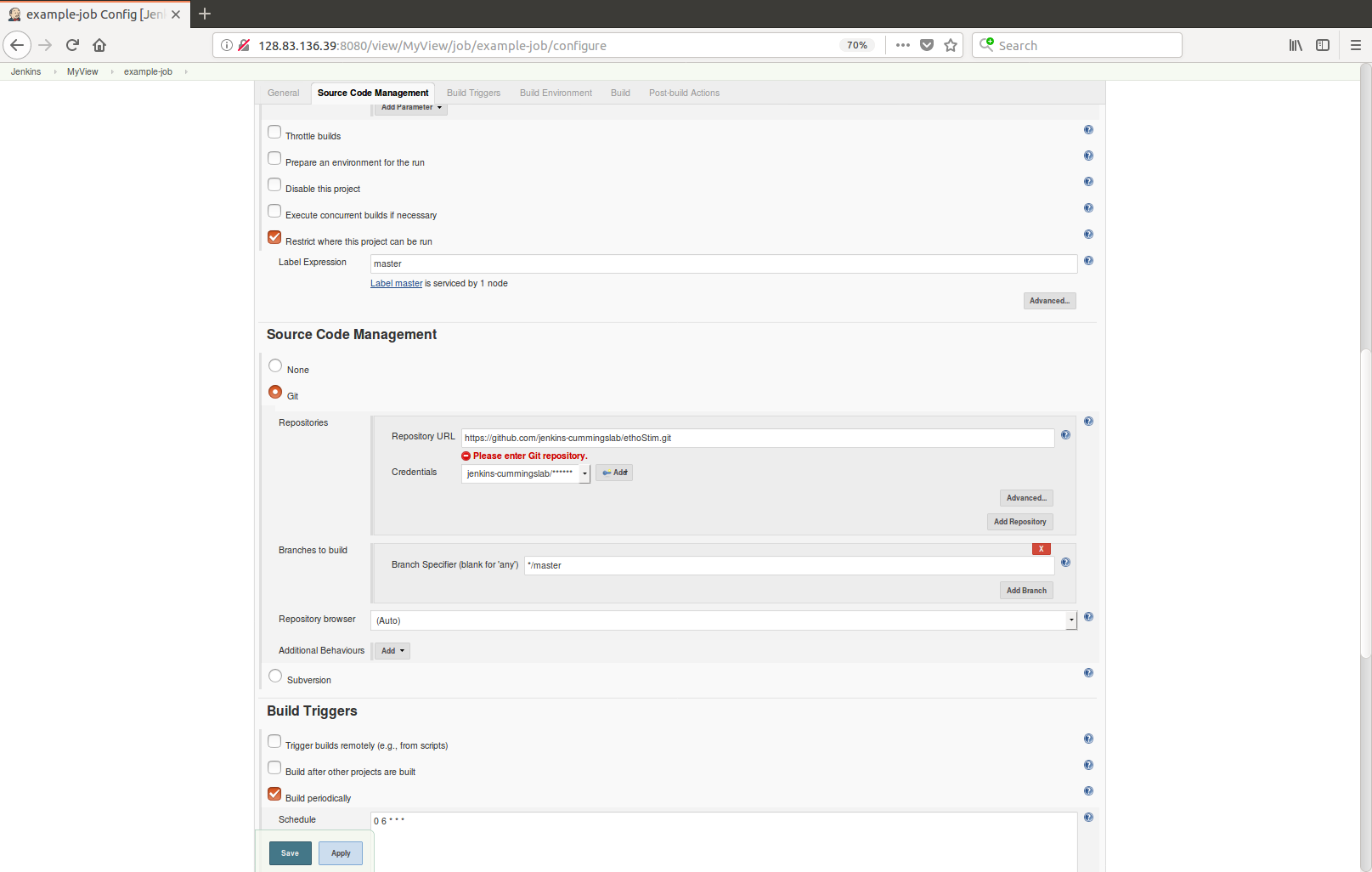

6. Configure Build Triggers settings such as periodic scheduling following a CRON-like scheduling format.

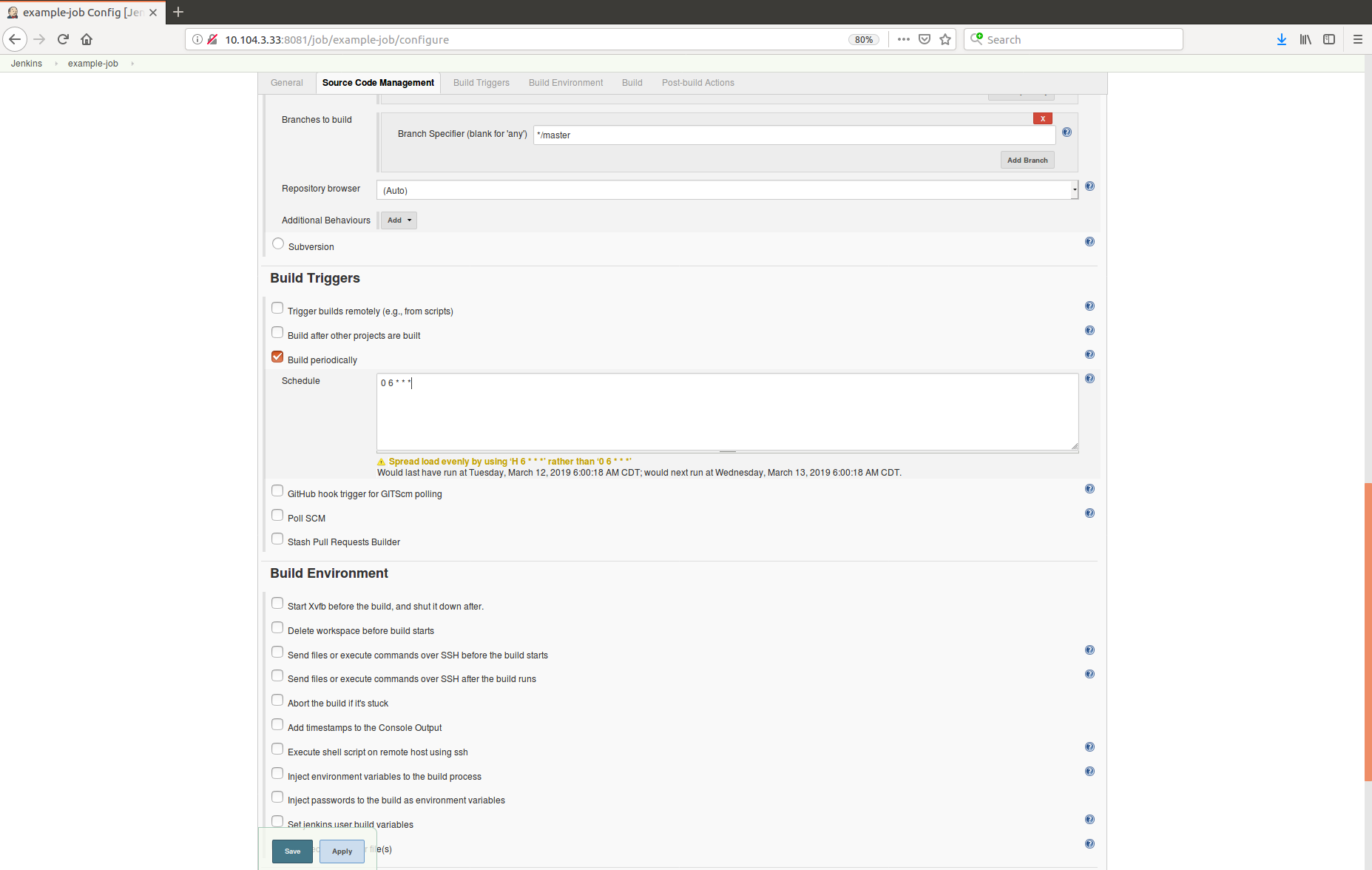

7. Configure Build Environment settings.

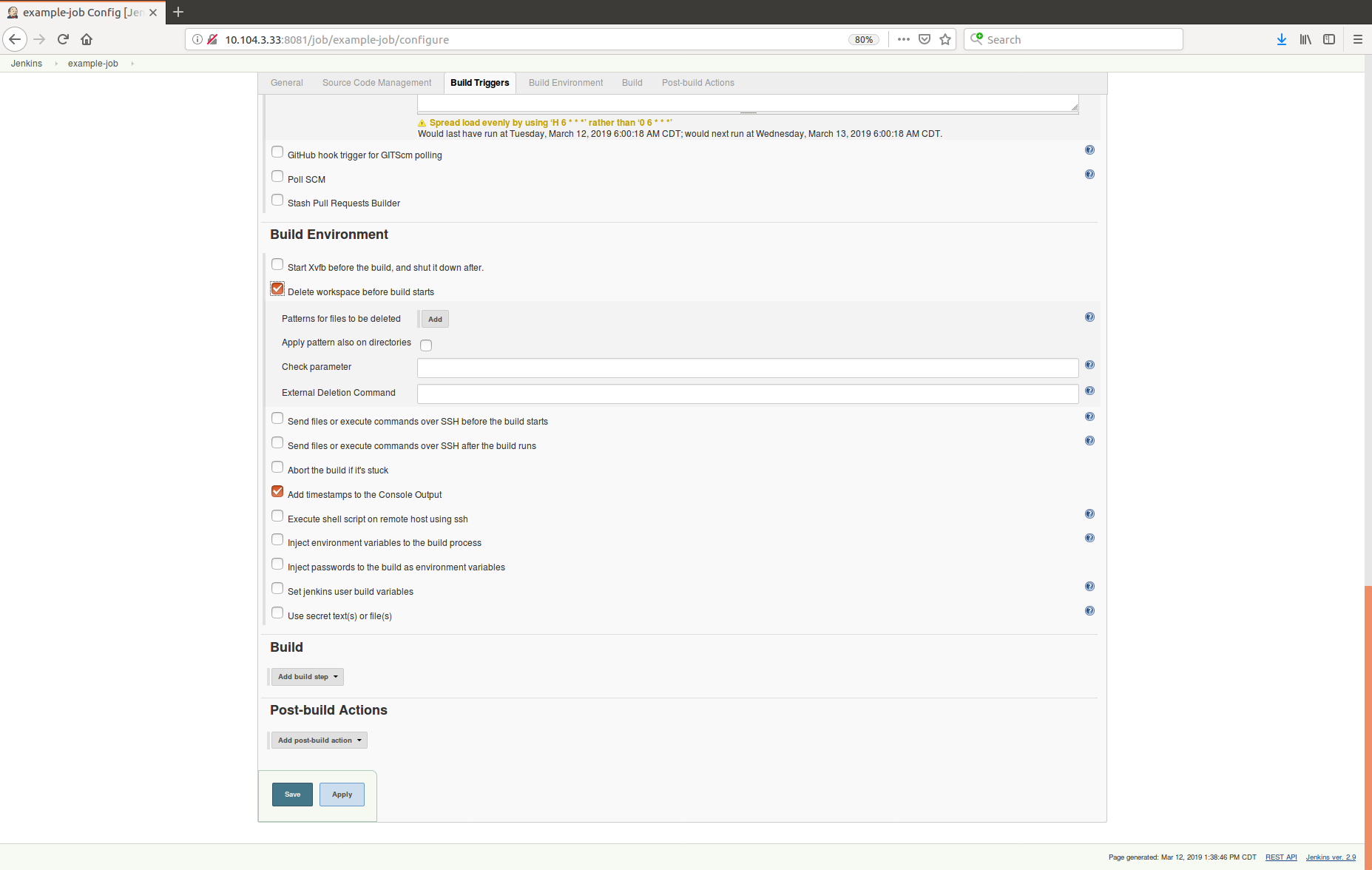

8. Configure the Build settings such as running a Python script from a Unix shell.

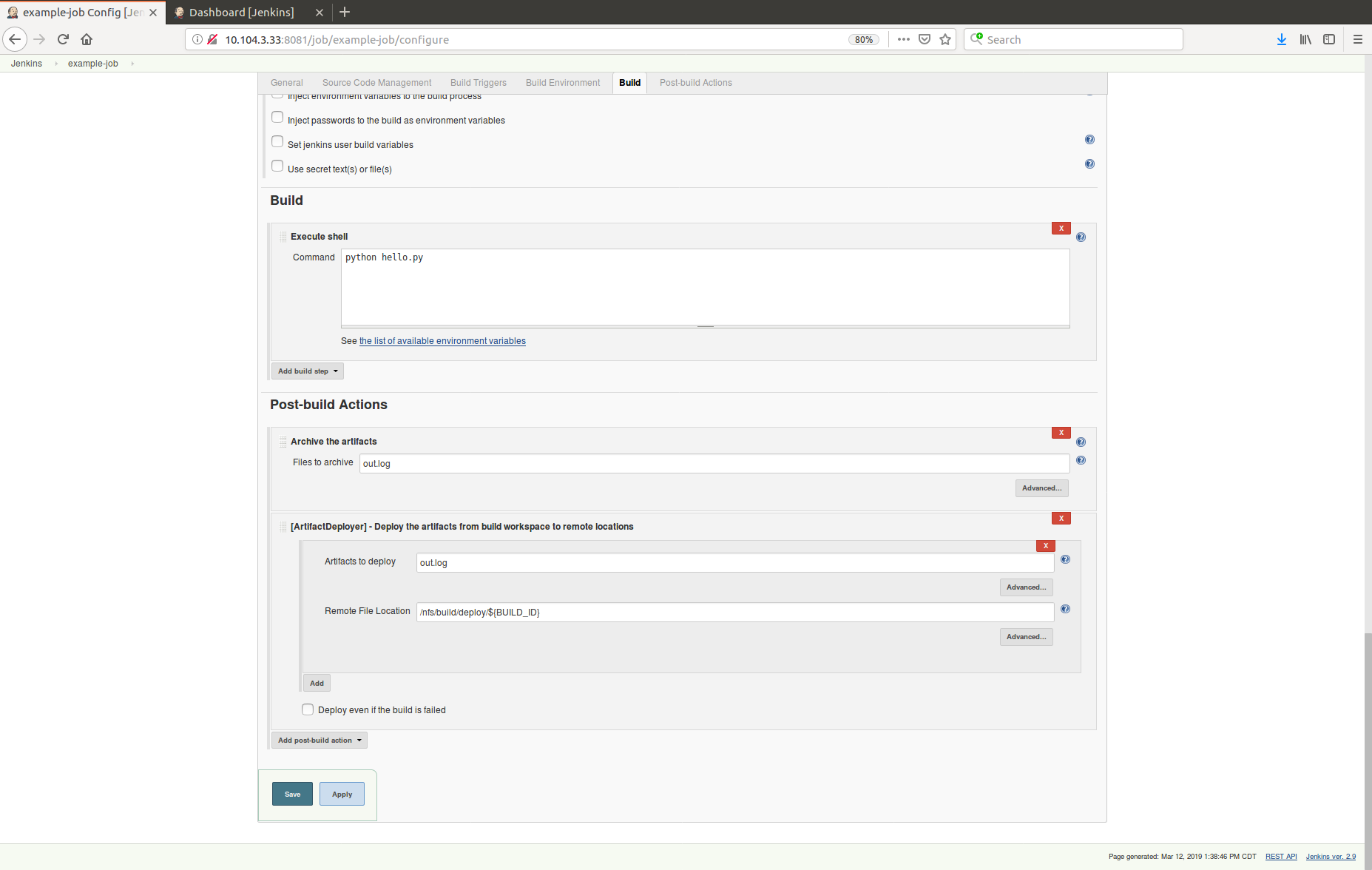

9. Configure the Post-build Actions such archiving log files and copying log files to a network share.

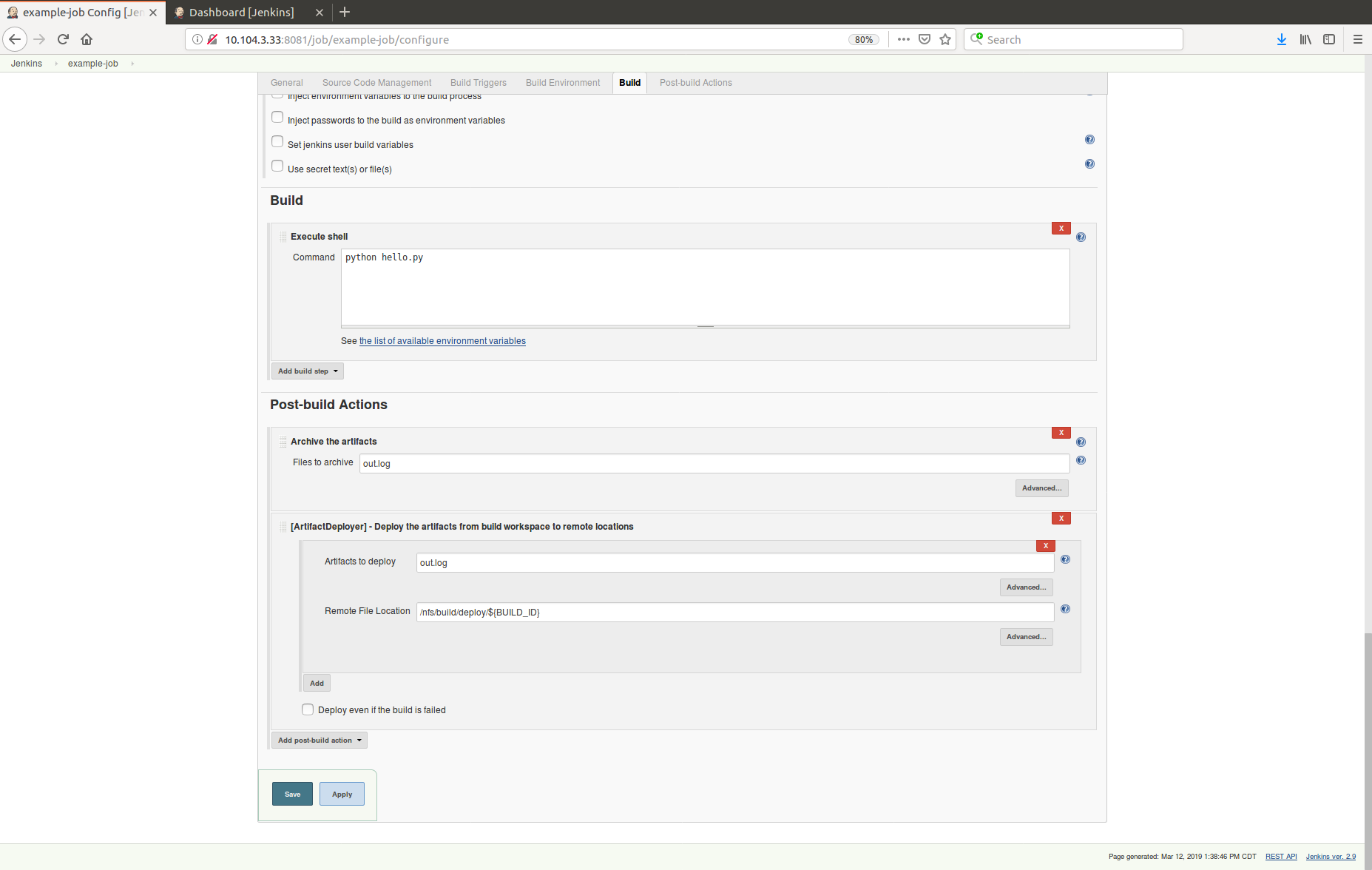

10. The new job will now run at the periodic interval or can be run immediately by clicking Build with Parameters, providing any configuration parameters, and clicking Build button.

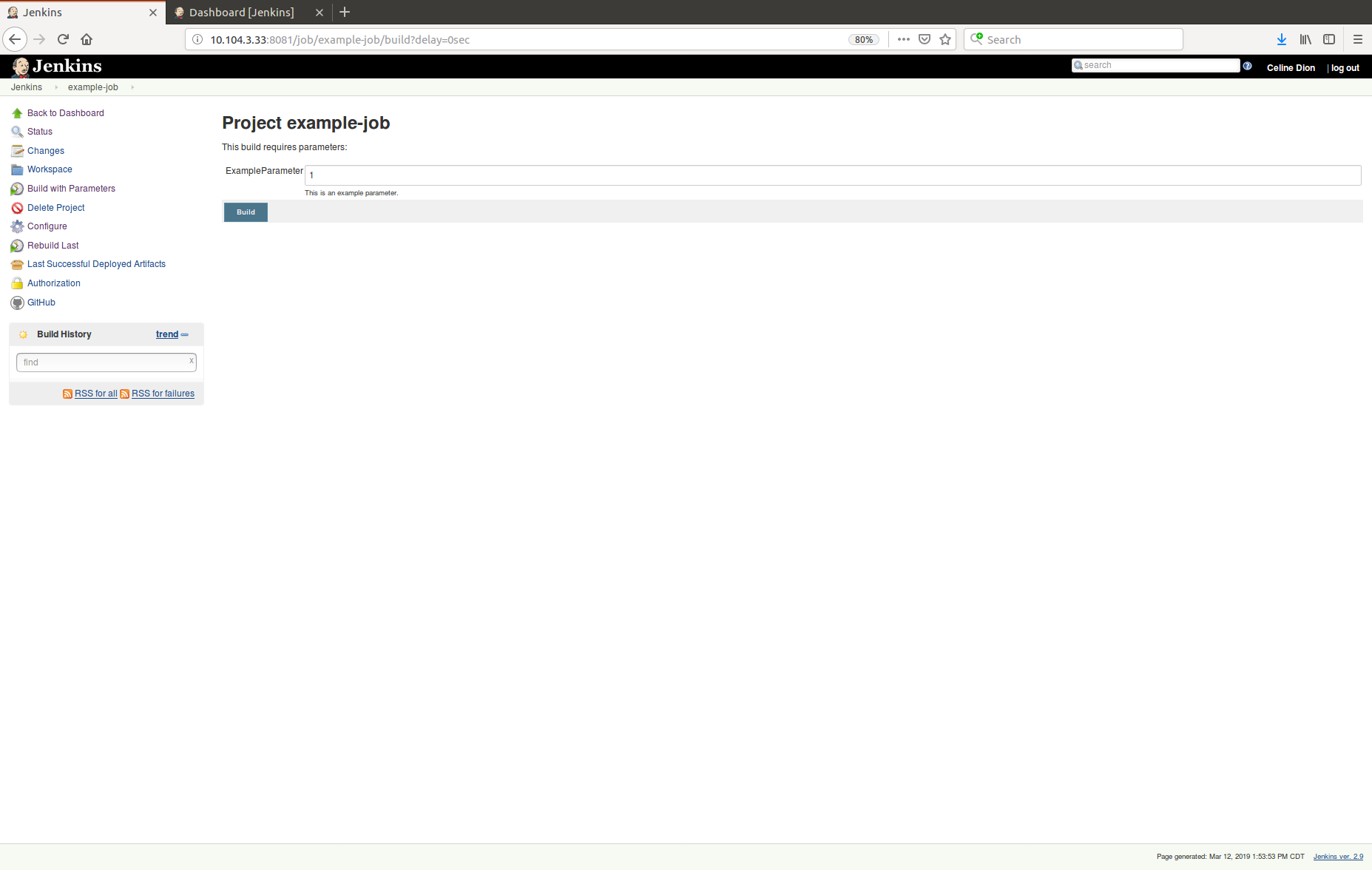

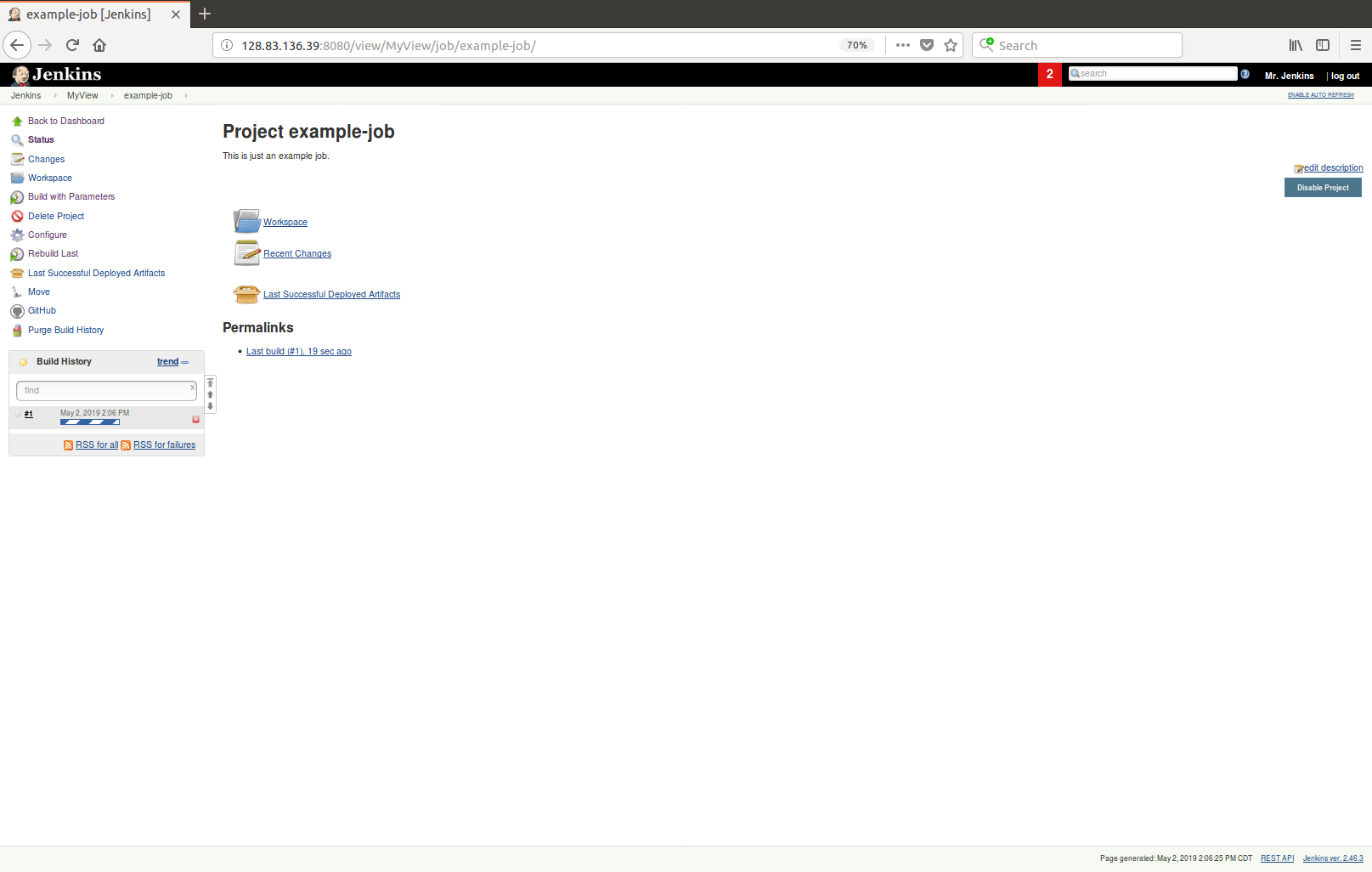

job is running!

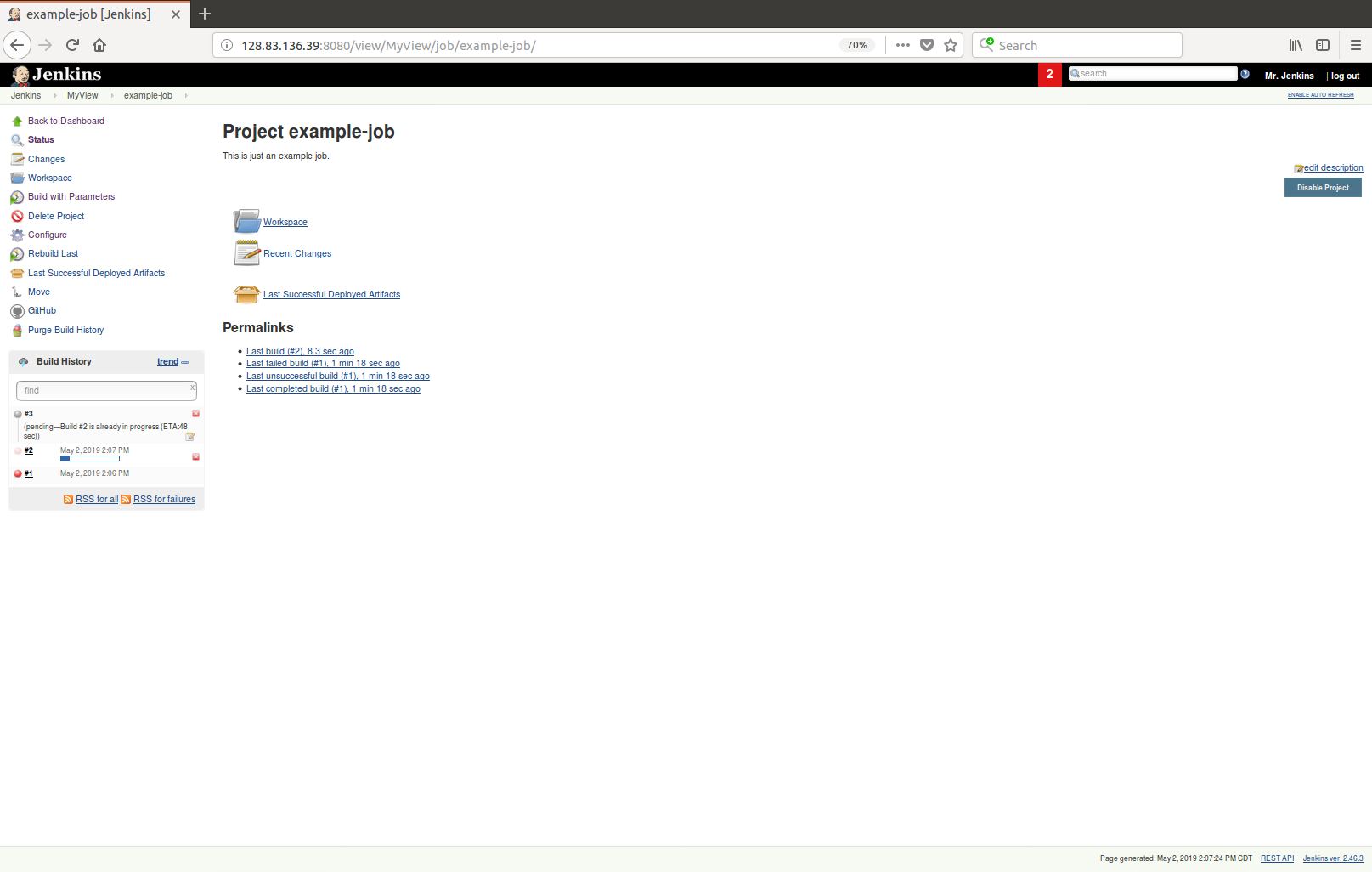

#1 completed (failed)

#2 is running

#3 is in the queue

The following figure highlights the Jenkins flow for our specific setup. A table follows that provides further detail of each Jenkins job. Jenkins jobs are setup to support round configuration, scheduling and execution of each trial, and cleanup. The central hub of the setup is a computer running Ubuntu referred as “alien”. In Jenkins terminology, this computer is configured as the “master node”. Each tank used in the experiment (up to 6 tanks per round) requires 2 RaspberryPi’s (1 per screen and feeder, camera is only connected to 1 of them). In Jenkins terminology, these computers are configured as “slave nodes”. Much of the heavy lifting and supporting functionality is executed from alien, but the actual trial script is executed on the RPis.

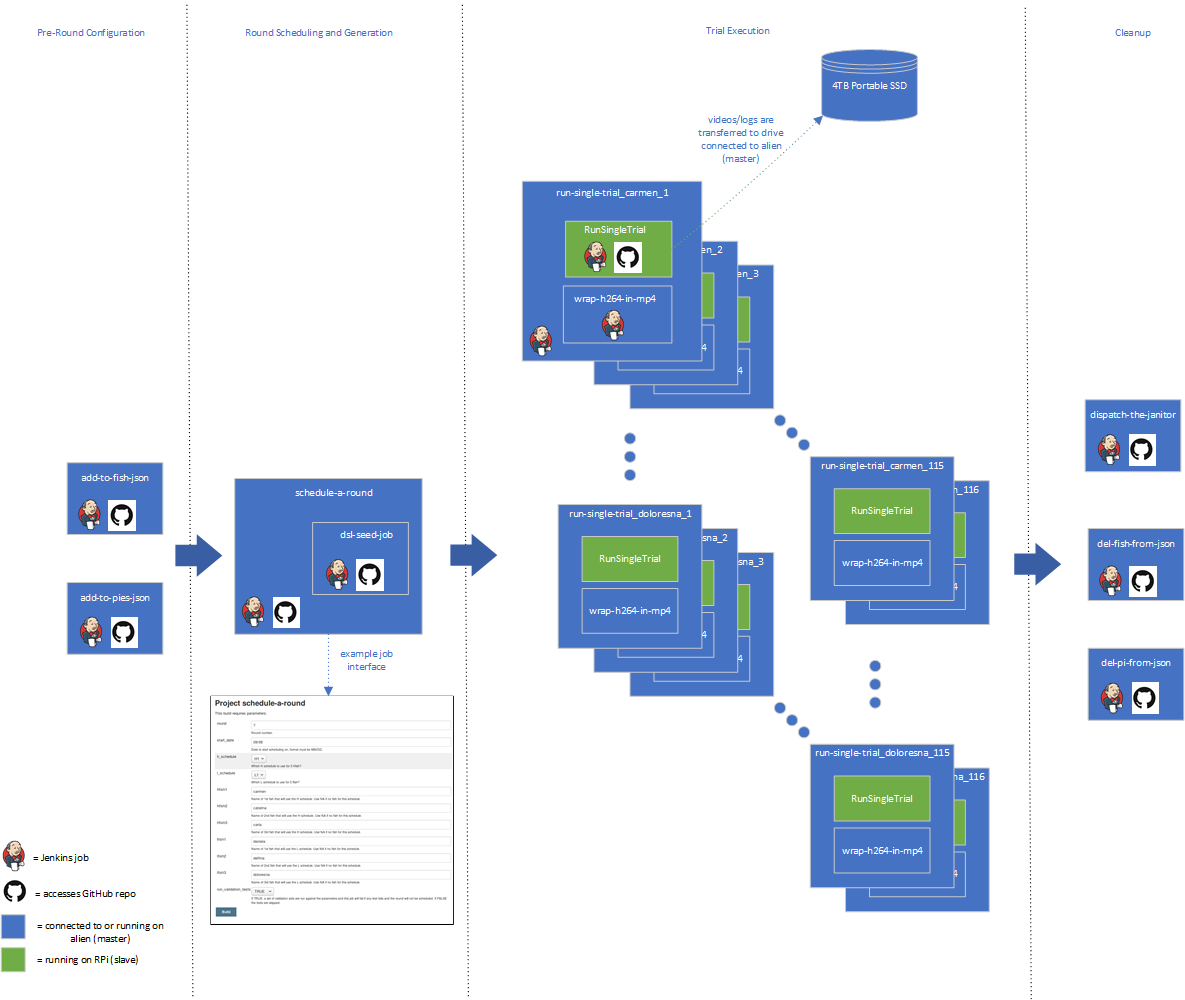

| Jenkins Job | Executing Node | Description | Scripts/Files^[2]^ |
| --- | --- | --- | --- |
| add-to-fish-json | alien (master) | Run during pre-round configuration to notify Jenkins about a fish and fish-specific parameters that will be used in testing. | add_to_fish_json.py  fish.json |
| add-to-pies-json | alien (master) | Run during pre-round configuration to notify Jenkins about a RPi and RPi-specific parameters that will be used in testing. | add_to_pies_json.py  pies.json |
| schedule-a-round | alien (master) | Run to configure and schedule a round of trials. User fills out a form specifying start date, schedule type, and up to 6 fish that will be tested. Optionally, user can also specify whether to validate trial parameters prior to round creation. This validation includes verifying that Jenkins “knows” all the fish, can communicate with all the applicable RPis, start date is not in the past or beyond 30 days in the future, etc. This job triggers another job, dsl-seed-job, that actually creates jobs to execute all the trials at the scheduled times. | mod_top_json.py  top.json  tests.py  tests.json |
| dsl-seed-job | alien (master) | Triggered by schedule-a-round. Based on round configuration, this job generates a separate job for each trial that will be scheduled based on the schedule type a given fish was configured for. The generated jobs use the following naming convention: run_single_trial_<fish>_<trial #>. | gen_trial_jobs.dsl  top.json  fish.json  pies.json  H#.json  L#.json |
| run_single_trial_<fish>_<trial #> | alien (master) | Container job created by dsl-seed-job that just triggers a RunSingleTrial job at a scheduled date/time with defined trial configuration parameters. At completion of RunSingleTrial, this job also triggers the job wrap_h264_in_mp4 before finalizing archived artifacts (i.e. video files, logs) on a centralized storage device. |  |
| RunSingleTrial | RPi (slave) | Pulls down echostim repository from GitHub and executes trial.py with provided parameters on a RPi. Once trial.py has successfully finished, the appropriate artifacts are moved off the RPI over the network to a centralized storage location. | trial.py  trial_config.json |
| wrap_h264_in_mp4 | alien (master) | This job utilizes the ffmpeg^[3]^ tool to wrap a h264 encoded video file into a mp4 container. This is done because more playback tools readily and fully support mp4. |  |
| del-fish-from-json | alien (master) | Clean-up job to remove a fish from Jenkins catalog. | del_fish_from_json.py |
| del-pi-from-json | alien (master) | Clean-up job to remove a RPi from Jenkins catalog. | del_pi_from_json.py |
| dispatch-the-janitor | alien (master) | Clean-up job to free up storage on RPis and other workspaces on the master node. | trashcan.json |

NOTE: It is possible to export a Jenkins configuration including jobs and import into another Jenkins setup. Contact one of the authors if interested in exploring.

1. If larger port switches are available, that would be preferred to reduce network latency.
2. These are scripts and files available in the ethoStim repository, <https://github.com/jenkins-cummingslab/ethoStim.git>
3. https://www.ffmpeg.org/
